# Coral reef potential connectivity in the southwest Indian Ocean

**DOI:** 10.1101/2023.11.23.568484

**Authors:** Noam S. Vogt-Vincent, April J. Burt, Rosa M. van der Ven, Helen L. Johnson

## Abstract

The tropical southwest Indian Ocean is a coral biodiversity hotspot, with remote reefs physically connected by larval dispersal through eddies and a complex set of equatorial and boundary currents. Based on multidecadal, 2 km resolution hydrodynamic and larval dispersal models that incorporate temporal variability in dispersal, we find that powerful zonal currents, current bifurcations, and geographic isolation act as leaky dispersal barriers, partitioning the southwest Indian Ocean into clusters of reefs that tend to consistently retain larvae, and therefore gene flow, over many generations. Whilst exceptionally remote, the Chagos Archipelago can broadcast (and receive) considerable numbers of larvae to (and from) reefs across the wider west Indian Ocean, most significantly exchanging larvae with the Inner Islands of Seychelles, but also the Mozambique Channel region. Considering multi-generational dispersal indicates that most coral populations in the southwest Indian Ocean are physically connected within a few hundred steps of dispersal. These results suggest that regional biogeography and population structure can be largely attributed to geologically recent patterns of larval dispersal, although some notable discrepancies indicate that palaeogeography and environmental suitability also play an important role. The model output and connectivity matrices are available in full, and will provide useful physical context to regional biogeography and connectivity studies, as well as supporting marine spatial planning efforts.

## 1 Introduction

Long-distance connectivity can be established between remote coral reefs through the process of larval dispersal, transported by ocean currents and modulated by larval behaviour (Cowen and Sponaugle, 2009). The physical transport of larvae between reefs is known as *potential connectivity* (Mitarai et al, 2009), and can be quantified as *explicit connectivity* (the likelihood of larval transport from *i* to *j*) or *implicit connectivity* (the likelihood of shared larval sources or destinations for *i* and *j*) (Ser-Giacomi et al, 2021). Physical larval transport can drive demographic and/or genetic connectivity, depending on the magnitude and variability of potential connectivity, and post-settlement processes (e.g. Watson et al, 2010; Lowe and Allendorf, 2010). Quantifying reef connectivity is therefore important for effective marine spatial planning (Beger et al, 2010; Balbar and Metaxas, 2019) and explaining biodiversity and biogeography (e.g. Cowen et al, 2006). However, these data are lacking in the tropical southwest Indian Ocean, a biodiversity hotspot (Obura, 2012; Veron et al, 2015; Kusumoto et al, 2020) home to around 7% of the world’s coral reefs (Souter et al, 2021).

At present, there is a basic understanding of reef connectivity in this region. The time-mean surface circulation in the southwest Indian Ocean is to first-order explained by Sverdrup transport, generating southward and northward interior transport north and south of *∼*15 *^◦^*S respectively (Godfrey, 1989). This drives the westward flowing South Equatorial Current (fig. 1). When the South Equatorial Current meets Madagascar, it splits into the Southeast and Northeast Madagascar Currents. The former flows southward along the east coast of Madagascar before feeding the Agulhas system, whilst the latter rushes past the north coast of Madagascar and forms a westward jet, generating eddies through barotropic instability in the process (Collins et al, 2014). The Northeast Madagascar Current then bifurcates, partially feeding the net-southward but eddy-dominated flow of the Mozambique Channel, but with most water entering the East African Coastal Current, which flows northward along the coast of Tanzania and Kenya (Swallow et al, 1991). The East African Coastal Current crosses the equator throughout much of the year (peaking during the southeast monsoon, in the austral winter), joining the Somali Current. During the northwest monsoon (austral summer), the Somali Current changes direction due to the monsoonal reversal of winds, meeting the East African Coastal Current south of the equator, and causing it to separate from the coast and feed the eastward South Equatorial Countercurrent (Schott and McCreary, 2001).

**Fig. 1.**
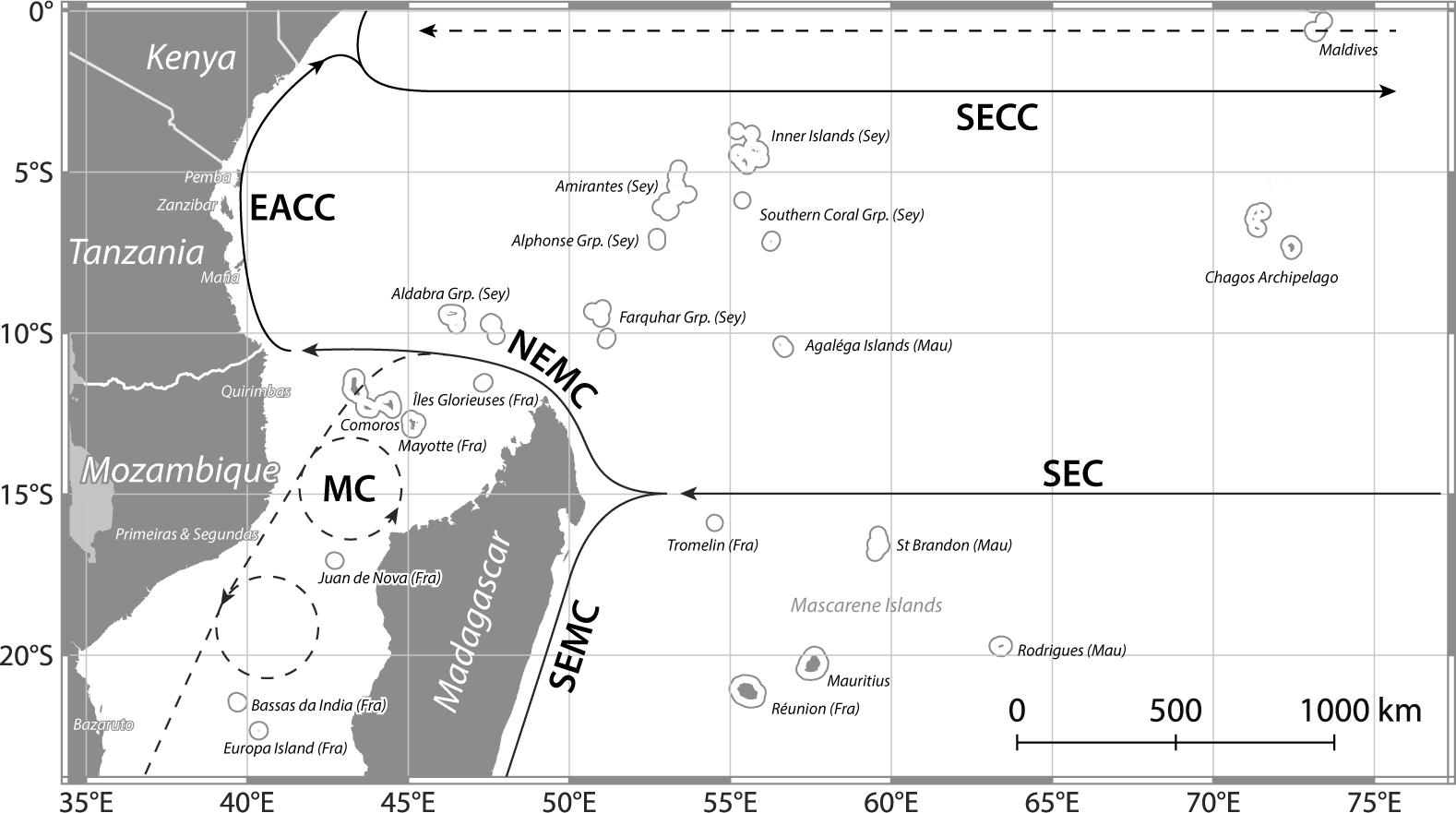
Map of the tropical southwest Indian Ocean (domain considered by this study), with a schematic representation of surface currents (during the northwest monsoon) based on Schott and McCreary (2001) (**SEC**: South Equatorial Current; **SECC**: South Equatorial Countercurrent; **NEMC**: Northeast Madagascar Current; **SEMC**: Southeast Madagascar Current; **EACC**: East African Coastal Current; **MC**: Mozambique Channel). Dashed lines represent transient surface currents. Note that small islands have been drawn with a halo for visibility.

There are limited data on broadcasting coral spawning in the tropical southwest Indian Ocean, with synchronised mass spawning taking place in September to December in Mozambique (Sola et al, 2016), whilst unsynchronised spawning has been observed or inferred elsewhere from October to April (Kenya, Mangubhai and Harrison, 2008a, 2009) and October to December (Seychelles, Koester et al, 2021). This indicates that most spawning takes place during the northwest monsoon, during which larval dispersal should be largely westward within the region dominated by the South Equatorial Current, with eddy-mediated dispersal across the Mozambique Channel, and strong connectivity along the path of the East African Coastal Current. This is broadly supported by regional coral biogeography (Obura, 2012), coral population genetics (e.g. van der Ven et al, 2022), and global (Wood et al, 2014) and regional (Crochelet et al, 2016; Mayorga-Adame et al, 2017; Gamoyo et al, 2019) larval dispersal simulations. These studies provide valuable insights into the connectivity of the southwest Indian Ocean, and generally support the existence of well-connected reef clusters in the northern Mozambique Channel and along the path of the East African Coastal Current.

However, since genetic connectivity is established over a large number of generations, the focus of these previous studies on snapshots of potential connectivity limits their ability to bridge the gap between numerical models and empirical observations (Legrand et al, 2022). Synthesising connectivity estimates across multiple metrics is essential for effective marine spatial planning (Balbar and Metaxas, 2019), which is becoming increasingly important given the growing anthropogenic stressors on coral reef systems in the southwest Indian Ocean (Obura et al, 2021). The importance of high-frequency variability for larval dispersal between remote reefs (Vogt-Vincent et al, 2023) also necessitates new approaches. The time-mean connectivity matrices considered by most previous studies neglect the considerable variability introduced by turbulence in the ocean, which has important implications for ecology (Watson et al, 2012). Finally, some genetic studies reveal complex, fine-scale patterns of genetic differentiation around small islands in the path of the EACC (e.g. van der Ven et al, 2016). These observations reveal variability below the scale that can be easily resolved by the large-scale larval dispersal studies conducted to date, which rely on relatively coarse oceanographic data (Wood et al, 2014; Crochelet et al, 2016; Gamoyo et al, 2019). Higher resolution hydrodynamic models may therefore improve our ability to discern the relative importance of physical larval dispersal (as opposed to post-settlement processes) in setting these localised patterns of differentiation.

The aim of this study is to describe and physically explain the potential connectivity between all shallow (*<* 20 m depth) reefs in the tropical southwest Indian Ocean, to support marine spatial planning efforts, and to test to what extent potential connectivity explains regional population structure and biogeography. For this purpose, we simulate daily coral spawning events based on 28 years of surface current predictions at 2 km resolution, representing the highest resolution dataset spanning the entire tropical southwest Indian Ocean, from East Africa to the Chagos Archipelago. In addition to the results presented here, all data from this study and scripts required to reproduce figures, as well as an interactive web app visualising the connectivity network, are available in the associated datasets (Vogt-Vincent et al, 2024).

## 2 Methods

### 2.1 Hydrodynamical and larval dispersal models

We modelled coral larvae as passively drifting particles, advected by surface currents, using the SECoW (Simulating Ecosystem Connectivity with WINDS) framework built on OceanParcels (Delandmeter and van Sebille, 2019), as previously described by Vogt-Vincent et al (2023). Young coral larvae are generally positively buoyant, but lose buoyancy with age. However, the vertical swimming speed of coral larvae is significant compared to vertical currents in the open ocean, and they therefore have some capacity to control their position in the water column (Szmant and Meadows, 2006). Larvae of at least some species appear to remain near the surface in response to pressure and light cues (Stake and Sammarco, 2003; Mulla et al, 2021), which we use to justify our assumption that coral larvae are confined to the upper water column.

We used surface currents from the WINDS model (Vogt-Vincent and Johnson, 2023), spanning the tropical southwest Indian Ocean from 1993-2020, at *∼*2 km spatial and 30-minute temporal resolution. Larval dispersal can take place over distances below this spatial resolution, due to asexual reproduction, short larval competency periods, and reef hydrodynamics (e.g. Grimaldi et al, 2022; Lord et al, 2023). It is therefore important to note that our analyses will only resolve dispersal over distances of at least a few kilometres. We released 2^10^ particles from each of the 8088 2 *×* 2 km coral reef cells identified in the tropical southwest Indian Ocean (Li et al, 2020) daily at midnight between October and March from 1993-2019, for a total of 4920 simulated spawning events. The available evidence suggests that coral spawning in the southwest Indian Ocean is concentrated in these months (e.g. Baird et al, 2021), but it is straightforward to repeat these analyses for alternative spawning seasonality, as the underlying dataset includes year-round spawning.

As in Vogt-Vincent et al (2023), the number of coral larvae represented by each particle is proportional to the surface area of reef within its origin cell (i.e. assuming constant fecundity per unit area of reef). We assumed that larvae gain and lose competency at constant rates following an initial pre-competency period, and die following a time-varying mortality rate (Connolly and Baird, 2010). Finally, we assumed that competent larvae settle with a constant probability per unit time (1 d*^−^*^1^), given that they are above a coral reef, thereby accounting for the possibility that larvae pass over a reef without settling (Hata et al, 2017). Since there is no meaningful information on the flow field below the resolution of the model grid (*∼*2 km), we scale this probability by the proportion of each grid cell that is occupied by reef. Further details on the larval dispersal model can be found in the supplementary materials (section 1). In this study, we focus on larval parameters for *Platygyra daedalea*, a broadcast spawning coral widespread across the Indo-Pacific and commonly used as a study taxon in the southwest Indian Ocean (e.g. Mangubhai et al, 2007; Mangubhai and Harrison, 2008b; Souter and Grahn, 2008; Montoya-Maya et al, 2016), with intermediate pelagic larval duration (Connolly and Baird, 2010). However, our first-order conclusions are robust across species (see supplementary materials, section 2). Analogous figures for the other sets of larval parameters described by Connolly and Baird (2010) can be found in the associated datasets.

### 2.2 Quantifying potential connectivity

For tractability, we first reduced the 8088 reef cells to 180 reef groups, identified using an agglomerative clustering algorithm. The definition of potential connectivity frequently used in the literature (e.g. Mitarai et al, 2009), i.e. the likelihood that a larva from reef *i* can physically settle at reef *j*, corresponds to the single-step *explicit* connectivity described by Ser-Giacomi et al (2021). We computed the long-term single-step explicit connectivity *C_ij_* from reef *i* to reef *j* as the time-mean proportion of larvae generated at reef *i* that settle at reef *j*. We similarly computed the standard deviation of explicit connectivity over time, *σ_ij_*. Although explicit connectivity is positively skewed and often dominated by extreme values, we display the mean explicit connectivity here rather than the median, as the latter is zero for most connections.

We also considered *implicit* connectivity, specifically the backward cumulated multi-step implicit connectivity (hereon CMIC), described by Ser-Giacomi et al (2021). In contrast to the single-step explicit connectivity, the CMIC considers ancestral larval sources over multiple steps of dispersal. For a pair of random walks respectively terminating at reefs *i* and *j*, CMIC*_ij_*(*k*) is defined as the probability that the pair of random walks passed through a common reef at least once within *k* steps. If we assume that gene flow through a reef network can be modelled as a random walk weighted by explicit connectivity (thereby neglecting all post-settlement processes), CMIC*_ij_*(*k*) may represent the degree of shared ancestry between two coral populations within *k* generations of dispersal. Although this assumption is unrealistic, Legrand et al (2022) nevertheless found that CMIC was a better predictor of genetic differentiation between populations than explicit connectivity alone. We estimated the number of generations of dispersal ‘separating’ two reefs, *G_ij_*, by solving for *k* such that CMIC*_ij_*(*k*) = 0.5.

Due to stochastic oceanographic variability, Vogt-Vincent et al (2023) found that CMIC based on time-varying explicit connectivity (i.e. allowing explicit connectivity to vary between spawning events) was only poorly represented by CMIC based on the time-mean explicit connectivity. We therefore computed *G_ij_* separately for 1000 randomly generated temporal subsets of the full time-varying explicit connectivity matrix (each subset representing a random series of spawning events, drawn from the 4920 available events), and took the median across this *G* ensemble.

### 2.3 Clustering

Partitioning the reef network into clusters that tend to retain coral larvae is a useful tool for understanding large-scale network structure, and comparing modelled connectivity to observations (e.g. Treml and Halpin, 2012; Thompson et al, 2018). For this purpose, we use the Infomap algorithm (Rosvall et al, 2009; Edler et al, 2023). Infomap partitions flow networks to minimise the information required to describe a random walk (e.g. gene flow through larval dispersal), and identifies clusters of nodes (reefs) that tend to retain flow (gene flow) more effectively than alternative metrics such as modularity (supplementary materials, section 3, Rosvall et al (2009)). However, due to the considerable stochastic variability in reef connectivity (Vogt-Vincent et al, 2023), clusters of reefs that tend to retain larvae based on time-mean potential connectivity may not be equivalent to clusters of reefs that tend to retain larvae over a given shorter time interval (thereby affecting short-term population dynamics). We are therefore interested in identifying clusters of reefs that *consistently* retain larvae over *l* spawning events.

Similar to our computation of *G* (section 2.2), we generated 1000 subsets of the full time-varying explicit connectivity matrix (each containing a random series of *l* spawning events), and computed the time-mean explicit connectivity for each subset over the *l* events. The value of *l* depends on the timescale of interest; we took *l* = 10 to investigate dispersal over decadal timescales, to reflect the response of reefs to short-term environmental disturbance. We then individually partitioned the reef network using Infomap for each short-term connectivity matrix, recording the lowest-level cluster assigned to each reef node in a cluster assignment matrix. A principal component analysis (PCA) revealed that over 80% of the variance in cluster assignment across the 1000 possible dispersal histories was explained by the first three principal components (PC1-3). We therefore reduced the cluster assignment matrix to three dimensions, with the closeness between reef nodes in this space representing how consistently reefs were assigned to the same cluster across stochastic oceanographic variability. We objectively assigned reefs to discrete clusters through K-Means clustering based on PC1-3. For illustrative purposes, we focus on the case with *k* = 8 clusters, as this distinguishes most clearly between the clusters emerging from the PCA. However, we note that the optimal number of clusters as identified by the elbow method appears to be around 3-5, depending on the pelagic larval duration (supplementary materials, fig. S1).

## 3 Results

### 3.1 Explicit connectivity

Fig. 2 shows the time-mean explicit connectivity matrix for *Platygyra daedalea*, which is also available as raw data, and presented as an interactive web application in the supplementary materials (Vogt-Vincent et al, 2024). Aside from the general pattern of time-mean explicit connectivity falling with distance, there is considerable asymmetry in the explicit connectivity, as expected from a region influenced by strong ocean currents (fig. 1). For instance, there is relatively strong explicit connectivity (*>* 10*^−^*^5^) from the Outer Islands of Seychelles to sites in East Africa due to the rapid Northeast Madagascar Current, whereas explicit connectivity in the opposite direction is orders of magnitude weaker as no direct dispersal pathway exists. The weakest connections within Seychelles are not the most distant (the Aldabra Group and Inner Islands, fig. 1), but rather from the Aldabra Group to the relatively nearby Farquhar Group, which would require larval transport in the opposite direction to the Northeast Madagascar Current. Asymmetric dispersal is also pronounced along the path of the northward East African Coastal Current from Tanzania to Kenya.

**Fig. 2.**
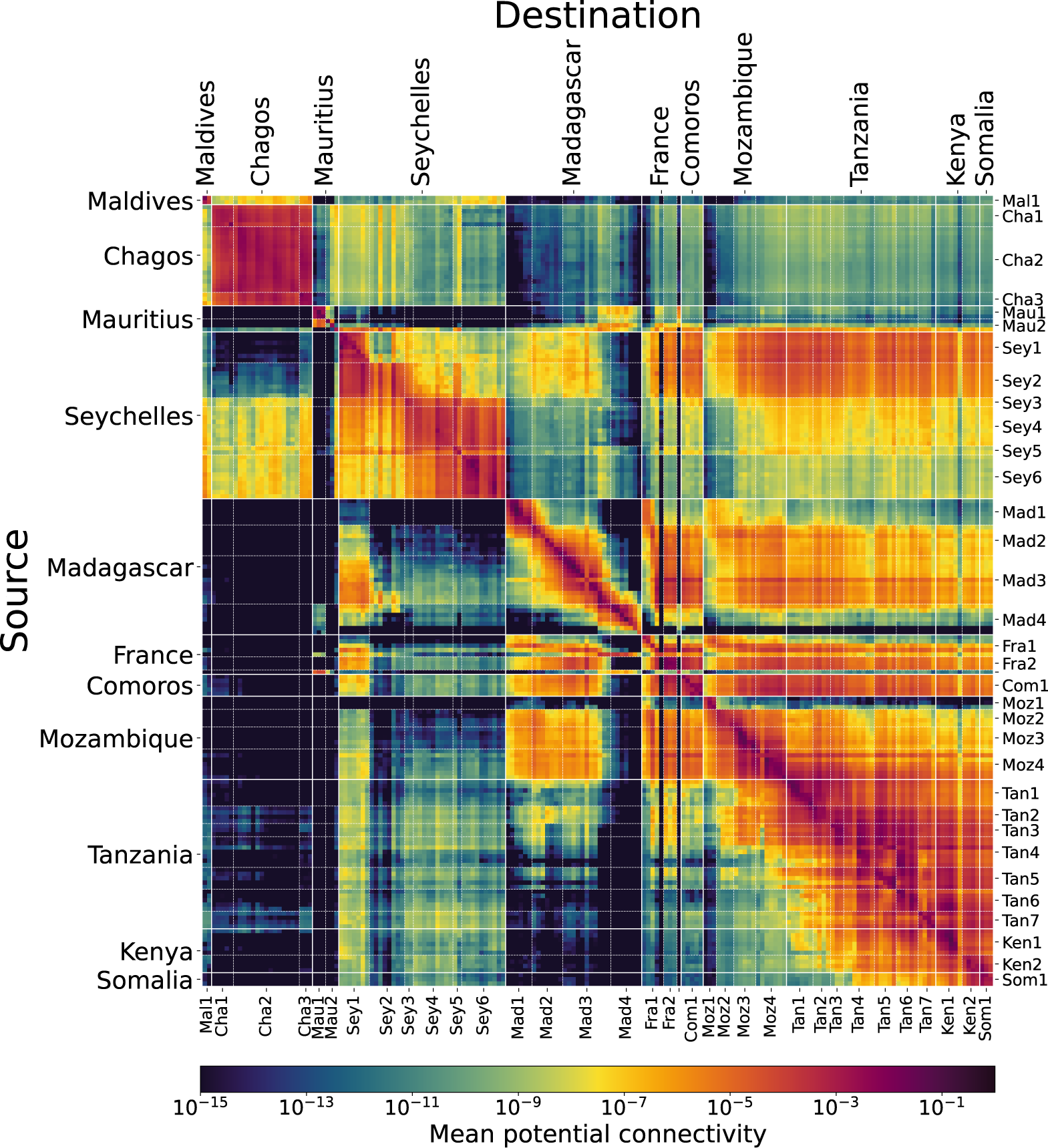
Time-mean explicit connectivity for *Platygyra daedalea* between pairs of reef groups. Subregion codes are as follows: **Mal1**: Maldives; **Cha1**: S Banks; **Cha2**: Grand Chagos Banks; **Cha3**: N Banks; **Mau1**: Mauritius (Island); **Mau2**: Outer Islands; **Sey1**: Aldabra Grp; **Sey2**: Farquhar Grp; **Sey3**: Alphonse Grp; **Sey4**: Amirantes; **Sey5**: Southern Coral Grp; **Sey6**: Inner Islands; **Mad1**: SW Madagascar; **Mad2**: NW Madagascar; **Mad3**: N Madagascar; **Mad4**: E Madagascar; **Fra1**: Scattered Islands; **Fra2**: Mayotte; **Fra3**: Réunion (not labelled); **Com1**: Comoros; **Moz1**: S Mozambique; **Moz2**: Primeiras & Segundas; **Moz3**: N Mozambique; **Moz4**: Quirimbas; **Tan1**: S Tanzania; **Tan2**: Songo Songo; **Tan3**: Mafia Island; **Tan4**: Dar es Salaam region; **Tan5**: Zanzibar; **Tan6**: Tanga; **Tan7**: Pemba; **Ken1**: S Kenya; **Ken2**: N Kenya; **Som1**: S Somalia. Please see supplementary materials, fig. S2 for a geographic key to subregion codes.

The strongest connection between the Chagos Archipelago and the rest of the southwest Indian Ocean is predicted to be with the Inner Islands of Seychelles, with mean explicit connectivity that can approach 10*^−^*^6^ from some islands in Seychelles, remarkable given the distance of over 1500 km. The eastward South Equatorial Countercurrent influences the Inner Islands of Seychelles during the northwest monsoon, providing an efficient pathway for eastward larval dispersal towards the Chagos Archipelago. Connections between the Chagos Archipelago and most other sites in the southwest Indian Ocean are expected to be principally in the opposite (westward) direction, following an initially slow southward pathway, before entering the South Equatorial Current and being rapidly transported towards the Mozambique Channel (fig. 1).

Explicit connectivity is consistently strong throughout the Chagos Archipelago, unsurprising given the expansive area of reef cover and relatively weak currents simulated by SECoW (minimising the proportion of larvae being lost to the open ocean). Larval dispersal is also strong within and between the constituent island groups of Seychelles. Due to the dominance of powerful mesoscale eddies within the Mozambique Channel, there is a fairly strong and symmetric exchange of larvae between west and north Madagascar and Mozambique, with explicit connectivity exceeding 10*^−^*^6^ between many sites.

The time-mean explicit connectivity matrix does, however, obfuscate the enormous temporal (and largely stochastic) variability in connectivity (supplementary materials, fig. S3). The standard deviation of explicit connectivity across daily spawning events is greater than the mean for practically all pairs of reef groups, with this ratio frequently exceeding 10, and even 100 for distant connections. Short-distance dispersal is generally associated with lower temporal variability, as is northward larval dispersal within Kenya and Tanzania due to the consistent flow of the East African Coastal Current, and westward dispersal from the Outer Islands of Seychelles, Comoros, and north Madagascar due to the Northeast Madagascar Current (Vogt-Vincent et al, 2023).

### 3.2 Implicit connectivity

Simplifying gene flow as a random walk weighted by explicit connectivity, fig. 3 (lower triangle) suggests that most pairs of *Platygyra daedalea* corals within the southwest Indian Ocean share a common ancestral population within around 200 steps (or generations) of dispersal. This is equivalent to around 1000 years, given that a typical broadcast-spawning coral reaches sexual maturity at an age of around 5 years (Rapuano et al, 2023). We expect particularly strong implicit connectivity within the Chagos Archipelago, the constituent island groups of Seychelles, and north and east Madagascar. Despite strong explicit connectivity, implicit connectivity along the path of the East African Coastal Current (Tanzania and Kenya) is relatively weak. The combination of strong currents and almost continuous reef cover along the coast results in larvae being distributed across a wide range of destination reefs, reducing the likelihood of any one individual connection (supplementary materials, fig. S4).

**Fig. 3.**
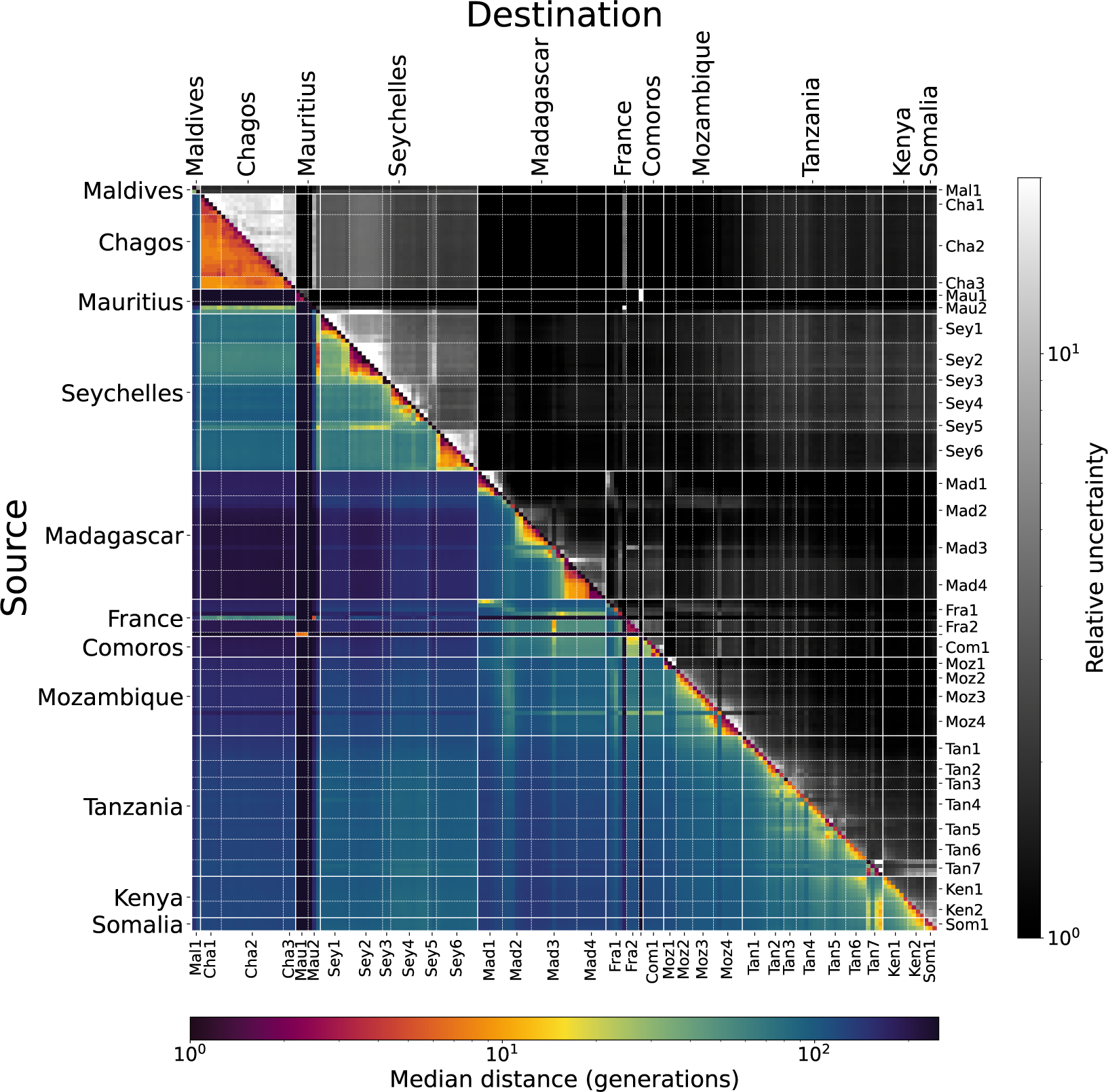
*Lower triangle*: Median distance between pairs of sites in generations, based on CMIC (*G_ij_*) across the 1000-member ensemble for *Platygyra daedalea*. *Upper triangle*: 95% confidence interval across the ensemble, divided by the median. Subregion codes as in fig. 2.

Our simulations predict that the implicit connectivity between the Chagos Archipelago and the rest of the southwest Indian Ocean is strongest with Seychelles, and three small island groups in the path of the South Equatorial Current (the Agaléga Islands, St Brandon, and Tromelin, fig. 1), all of which physically share ancestral populations within 10-100 dispersal steps for *Platygyra daedalea*. This underlines the potential importance of Seychelles in maintaining gene flow across the Indian Ocean by acting as a stepping-stone for dispersal.

A small subset of reefs are considerably more isolated in terms of implicit connectivity, namely (the island of) Mauritius, Rodrigues, and Réunion. For corals with similar larval parameters to *Platygyra daedalea*, these reefs are separated by over 200 steps of dispersal from practically all other reef sites in the southwest Indian Ocean. This is not obvious from the single-step explicit connectivity matrix, since we expect that these reefs all send considerable numbers of larvae to Madagascar. This dispersal is strongly unidirectional, which may explain why the implicit connectivity (shared ancestry) is so low.

In contrast to single-step explicit connectivity (section 3.1), variability in multistep implicit connectivity reduces with distance (fig. 3, upper triangle). For instance, although our simulations predict that pairs of reefs within the Chagos Archipelago are implicitly separated by around 10 steps of dispersal, this can vary by more than an order of magnitude depending on when exactly a spawning event takes place. The larger number of steps of dispersal required to connect distant pairs of reefs instead smooths out variability and increases redundancy, so the relative uncertainty in implicit connectivity between distant sites may, in fact, be lower than for nearby sites. This is a positive finding for large-scale biogeographic or population genetic studies, as we may therefore expect greater agreement between model predictions and observations over larger spatial scales (as compared to more localised studies).

### 3.3 Clustering

Infomap identifies a number of clusters of reefs that consistently retain larval flow despite variability due to dynamic ocean currents (fig. 4), which we call *meta-clusters*.

**Fig. 4.**
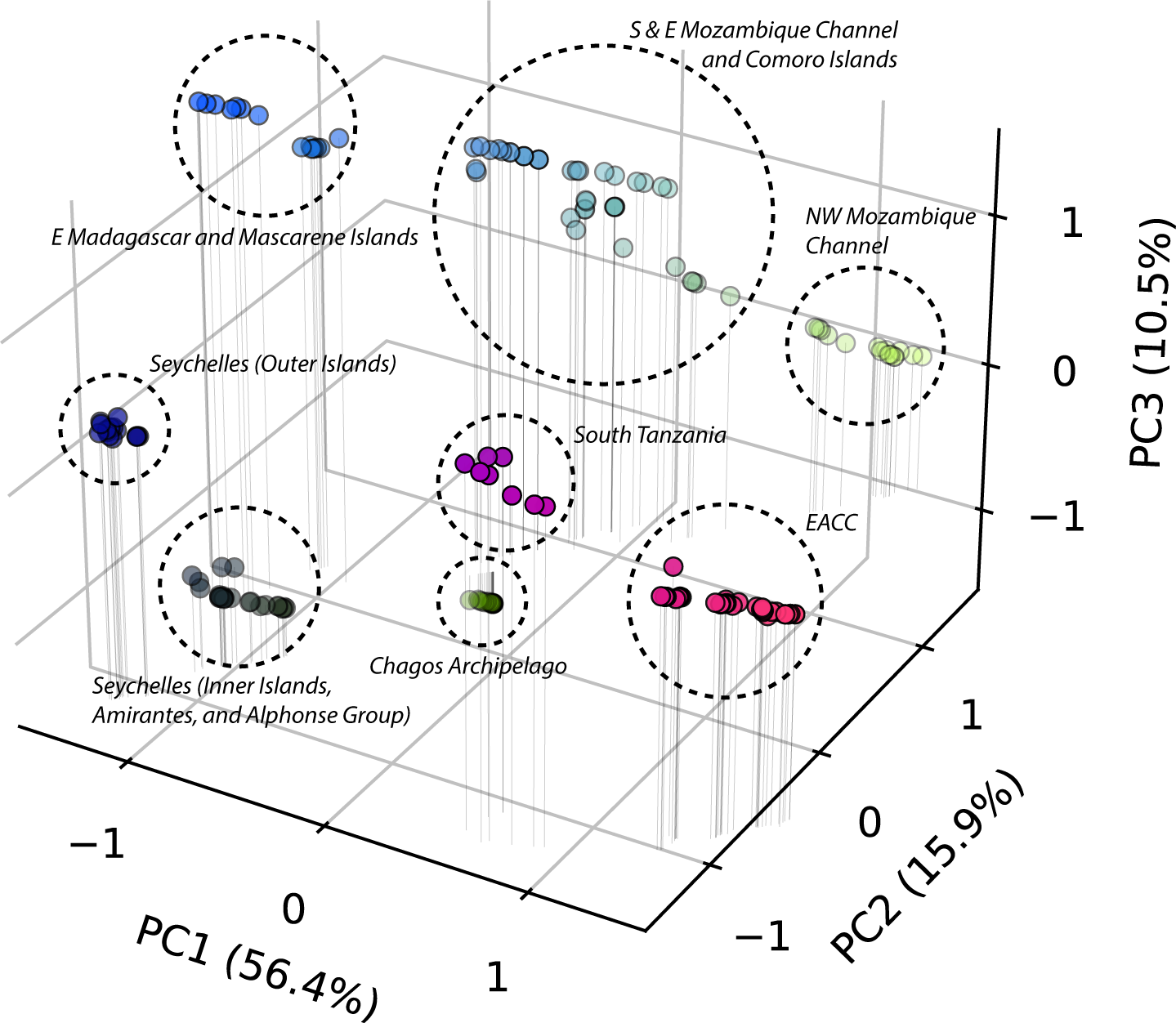
The 180 reef groups (for *Platygyra daedalea*) plotted according to their first three principal components. The normalised values of PC1, PC2, and PC3 respectively also determine the red, green, and blue values of each group. The meta-clusters identified by dashed lines are based on K-Means clustering with 8 clusters, see supplementary materials, figs. S7-8 for an equivalent plot with discrete colouring by *k* = 3 and *k* = 8 K-Means clusters. Note that the principal components are normalised in this figure for clarity, but the true Euclidean distance between groups is plotted in the supplementary materials, fig. S5.

Some meta-clusters are particularly distinct, attributable to geographic isolation and/or weaker currents (such as the Outer Islands of Seychelles and the Chagos Archipelago). Other reefs appear to plot across more of a continuum, such as Madagascar and the broader Mozambique Channel region. Computing the Euclidean distance between reef groups in principal component space (supplementary materials, fig. S5) therefore represents a distinct similarity metric to the implicit connectivity, representing how consistently reefs are assigned to clusters that retain larval flow.

By assigning the values of the three principal components to red (PC1), blue (PC2), and green (PC3) channels respectively, we can plot meta-clusters geographically (fig. 5). We note that some caution is needed when interpreting this figure; the three principal components are not equally important, and RGB colour space is not perceptually uniform (Thyng et al, 2016). Nevertheless, fig. 5 represents an intuitive way to visualise the opacity of dispersal barriers (represented by sharp colour and brightness changes), and their position relative to geographic and oceanographic features.

**Fig. 5.**
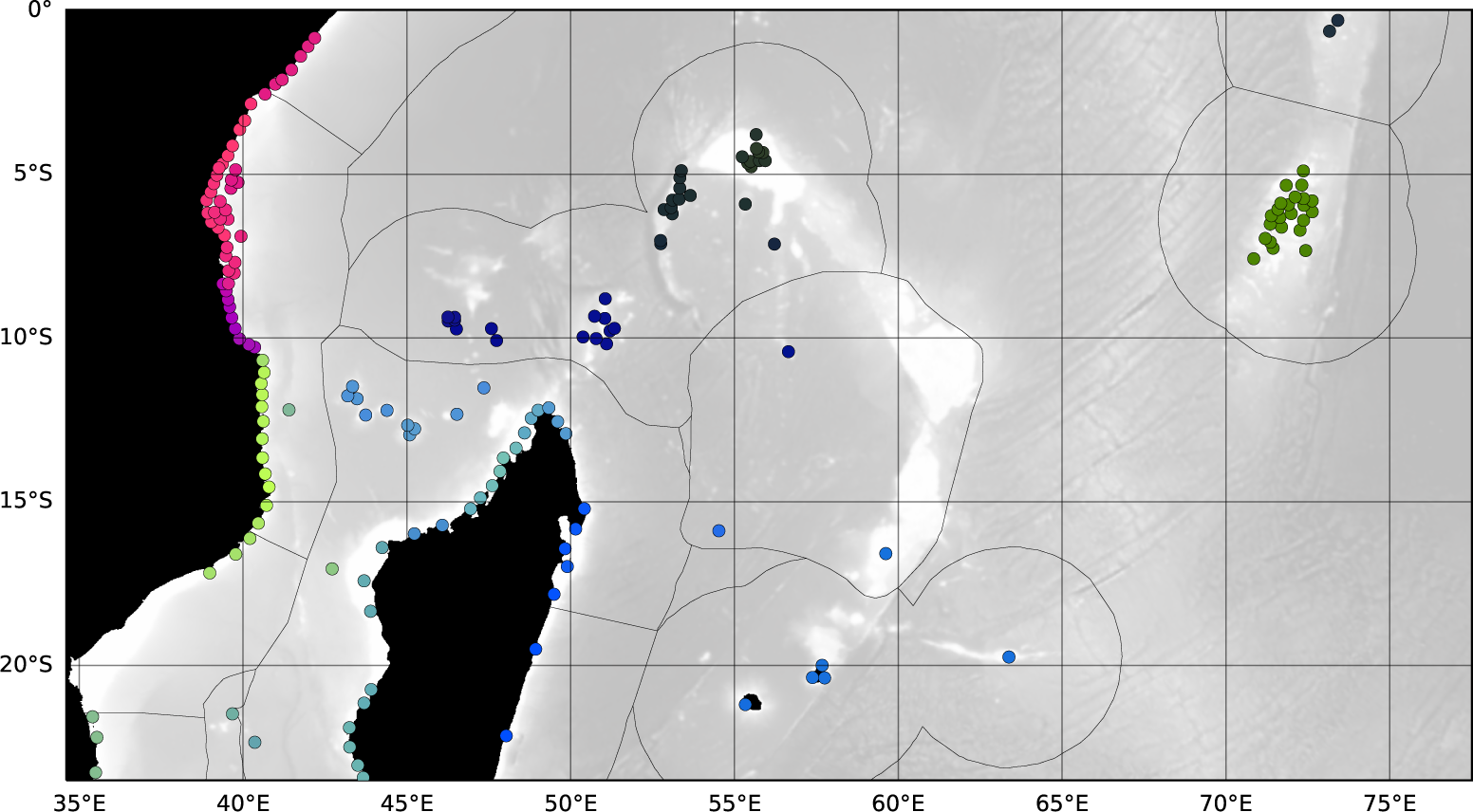
Reef groups plotted geographically, coloured according to their position in principal component space, based on larval parameters for *Platygyra daedalea* (fig. 4). Black lines represent the boundaries of general biogeographical ecoregions (Spalding et al, 2007), and bathymetry (GEBCO Compilation Group, 2022) is plotted for reference in the background. See supplementary materials, figs. S7-8 for equivalent plots with discrete colouring by *k* = 3 and *k* = 8 K-means clusters.

## 4 Discussion

### 4.1 Physical barriers to gene flow

Divisions between meta-clusters (consistent barriers to dispersal) tend to clearly coincide with persistent oceanographic and geographic barriers. For instance, the strong westward flow of the Northeast Madagascar Current limits the potential for northsouth larval exchange between the Outer Islands of Seychelles and the Mozambique Channel region, despite their geographic proximity. The westward flow of the South Equatorial Current similarly prevents larval dispersal between Seychelles and the Mascarene Islands, although the geographic isolation is certainly also important.

The bifurcations of the South Equatorial Current into the Northeast and Southeast Madagascar Currents, and the Northeast Madagascar Current into the East African Coastal Current and net-southward flow within the Mozambique Channel (fig. 1) result in consistent dispersal barriers in our simulations (also identified in previous simulations by Crochelet et al (2016); Gamoyo et al (2019)). Less obviously, a minor barrier appears in our meta-clustering analysis between south and north Tanzania, around Mafia Island (fig. 4). Mafia Island deflects the path of the East African Coastal Current, potentially sheltering reefs in its wake from upstream larvae (supplementary materials, fig. S6) and thereby generating a (leaky) break in connectivity along the coast of Tanzania. This is also visible as a slight discontinuity in the time-mean explicit connectivity matrix (fig. 2).

Some physical dispersal barriers only appear using the multi-generational connectivity metrics that consider higher-order network structure. In our meta-clustering analysis, the Comoro Islands tend to cluster with north Madagascar rather than Mozambique. Although eddies generated in the wake of north Madagascar provide a dispersal pathway for larvae from the Comoro Islands to Madagascar, larvae are nevertheless more likely to reach reefs in Mozambique, particularly the northern coast and Quirimbas Archipelago (fig. 2). However, over many steps of dispersal, a random walk is more likely to remain within the Comoros-Madagascar system than the Comoros-Mozambique system (which is also apparent in the CMIC, fig. 3), suggesting that the barrier to gene flow between Comoros and Mozambique may be stronger in the long term.

Similarly, although larvae leaving the Aldabra Group in Seychelles are considerably more likely to reach East Africa than the Inner Islands of Seychelles (due to the Northeast Madagascar Current), our meta-clustering analysis suggests that gene flow tends to be retained within the Seychelles system (i.e. we may expect greater similarities between corals of Aldabra and the other Outer Islands of Seychelles, than with corals in East Africa). This may be due to strong multi-generational gene flow from the Inner Islands of Seychelles to the Aldabra Group via the Amirante Islands (fig. 2) overwhelming the larval ‘leakage’ from the Aldabra Group.

The three principal components identified by the meta-clustering analysis (fig. 4) individually have relatively clear physical interpretations (supplementary materials, fig. S9-11). The dominant principal component (PC1) is high in reefs influenced by the East African Coastal Current, and low elsewhere. The East African Coastal Current is therefore arguably one of the most important hydrographic features in the southwest Indian Ocean for coral reef connectivity, as the strength of the mean flow facilitates consistent, strong reef connectivity along its path. PC2 is highest within the Mozambique Channel and Chagos Archipelago. This represents the important role of eddies and boundary currents within the Mozambique Channel in facilitating coral reef connectivity, as well as the role of the South Equatorial Current in transporting larvae from the Chagos Archipelago towards the Mozambique Channel region. Finally, PC3 is elevated east of Madagascar, across the Comoro Islands, and along the path of the East African Coastal Current, representing connectivity between the South Equatorial Current and the East African Coastal Current. It is not surprising that these last two principal components represent the importance of downstream linkages from the South Equatorial Current. Surface waters in the East African Coastal Current are largely sourced from the South Equatorial Current (Swallow et al, 1991), and variability in Mozambique Channel transport is closely tied to variability in the South Equatorial Current (Ridderinkhof et al, 2010; Backeberg and Reason, 2010). Indeed, ‘switching’ in downstream connectivity from the South Equatorial Current to East Africa versus the Mozambique Channel were two major modes of low-frequency reef connectivity variability identified by Vogt-Vincent et al (2023), using the same SECoW model as the present study.

These major dispersal barriers are consistent across the five larval species described by Connolly and Baird (2010), although the relative importance of some barriers vary. In general, meta-clusters become increasingly fragmented for larvae spending less time adrift, in agreement with previous studies (e.g. Thomas et al, 2014; Gamoyo et al, 2019). For corals with the shortest pelagic larval duration, this fragmentation introduces new dispersal barriers, such as within west Madagascar. Here, fragmented reef habitat, complex coastline, and coastal currents favouring retention during spawning months (Vogt-Vincent et al, 2023) may disproportionately affect short-lived larvae. Unsurprisingly, the Chagos Archipelago is significantly more isolated with a shorter pelagic larval duration, which particularly affects connectivity with the northern Mozambique Channel. Conversely, connectivity between the Inner and Outer Islands of Seychelles is proportionately stronger for short-lived larvae, suggesting that Seychelles might become an increasingly distinct biogeographic unit for these species. The dispersal barrier due to the bifurcation of the Northeast and Southeast Madagascar Current also appears to be relatively less important for corals with a short pelagic larval duration, perhaps due to the proportionately greater reef network fragmentation elsewhere in Madagascar. Further details are available in the supplementary materials (section 2).

### 4.2 The role of dispersal in shaping biogeography and population structure

Although local adaptation can result in fine-scale phenotypic and genetic differentiation below the scale of dispersal (e.g. Sanford and Kelly, 2011; Hays et al, 2021), larval dispersal nevertheless plays an important role in setting population structure in the marine environment (e.g. Carpenter et al, 2011; Pinsky et al, 2017; Legrand et al, 2022). Comparison of the meta-clusters derived from our analyses with marine ecoregions identified by Spalding et al (2007) and Obura (2012) demonstrates that there is considerable agreement between clusters of reefs that consistently retain coral larvae, and regional biogeography across a range of taxa, including reef-building corals (fig. 5). In particular, these ecoregions reflect the role of the Northeast Madagascar Current as a dispersal barrier, the strength of connectivity within Seychelles, the clustering of the Comoro Islands with Madagascar, strong gene flow along the path of the East African Coastal Current, and the existence of significant genetic connectivity between the Chagos Archipelago and the rest of the southwest Indian Ocean (Obura (2012) specifically).

Obura (2012) proposed a ‘Northern Mozambique Channel’ ecoregion (characterised by high biodiversity and similar coral fauna), and noted that this ecoregion also shared considerable similarity with corals found in the Chagos Archipelago. For coral species with a moderate-to-high pelagic larval duration, K-Means clustering suggests the optimum number of meta-clusters is 3. One of these meta-clusters closely corresponds to this proposed ecoregion, containing the entire Mozambique Channel region and the Chagos Archipelago (supplementary materials, fig. S7). This grouping is not identical to the ecoregion suggested by Obura (2012) (for instance, the northern boundary is at the border between Mozambique and Tanzania, rather than at Mafia Island). Nevertheless, this represents strong evidence that the observed faunal similarities between the Chagos Archipelago and Mozambique Channel identified by Obura (2012) are grounded in the geography of larval dispersal. However, although Obura (2012) suggested that the South Equatorial Countercurrent (fig. 1) could play a role in connecting the Chagos Archipelago with the Northern Mozambique Channel ecoregion, our simulations suggest that this dispersal pathway is negligible (fig. 2). Instead, any direct physical connectivity between the Chagos Archipelago and the Mozambique Channel is likely unidirectional and westward, through the South Equatorial Current.

The tendency of our analyses to group the entire Mozambique Channel region together is indicative of frequent larval dispersal across the basin, further supported by population genetics from a broadcasting coral (van der Ven et al, 2022). However, with finer granularity, a (permeable) dispersal barrier emerges across the Mozambique Channel (fig. 5), placed by our analysis to the west of the Comoro Islands, agreeing well with the location of an ecoregion boundary proposed by Spalding et al (2007). van der Ven et al (2021) identified greater genetic differentiation across this barrier in the northern Mozambique Channel for a brooding coral, indicating that this dispersal barrier may be particularly important for corals with a lower pelagic larval duration (although stronger connectivity was maintained between Mozambique and southwest Madagascar).

In contrast to our predictions, neither Spalding et al (2007) nor Obura (2012) identified an ecoregion boundary at the border between Mozambique and Tanzania (*∼* 10*^◦^*S). There is also no strong support for a barrier here from coral population genetics (van der Ven et al, 2021, 2022), although a barrier was identified at a similar location in previous modelling study (Gamoyo et al, 2019). Instead, Obura (2012) placed an aforementioned ecoregion boundary at Mafia island (*∼* 7.5*^◦^*S). This is consistent with the potential role, as identified by our simulations, of flow diversion around Mafia Island acting as a barrier to larval dispersal, and is potentially supported (albeit equivocally) by some genetic studies (van der Ven et al, 2016, 2021). On the other hand, no such barrier was identified by Spalding et al (2007), or other population genetic analyses (e.g. Souter and Grahn, 2008; van der Ven et al, 2022). Spalding et al (2007) instead identified an ecoregion boundary in northern Kenya. There is some evidence for a dispersal barrier here in our simulations for corals with a shorter pelagic larval duration (supplementary materials, section 2; fig. S12), with this barrier being generated by the confluence of the East African Coastal Current and Somali Current during the northwest monsoon.

In general, however, the prediction of strong coral connectivity along the path of the East African Coastal Current is strongly supported by coral population genetics (Souter and Grahn, 2008; Souter et al, 2009; van der Ven et al, 2016, 2021, 2022), although there is also evidence for local-scale genetic differentiation. For instance, Souter et al (2009) found significant differentiation between corals sampled in northeast and west Zanzibar, and most other samples in the region. van der Ven et al (2016) also identified a distinct cluster within the Pemba and Zanzibar channels. A possible explanation for the isolation of corals in northeast Zanzibar is the trapping of larvae along the coast due to the deflection of the EACC as it passes the southern tip of the island (Vogt-Vincent et al, 2023). Due to comparatively sluggish flow within the Zanzibar Channel, most larvae generated at sites in west Zanzibar remain within the Zanzibar Channel rather than entering the EACC, again resulting in self-recruitment (also found by Mayorga-Adame et al (2017)). However, this cannot explain why a separate, nearby site in west Zanzibar showed little differentiation from other reefs in the region (Souter et al, 2009). This is an important reminder that, although large scale biogeography and population genetics may be strongly grounded in regional scale oceanography, reef scale hydrodynamics and environmental variability which are not captured by our simulations may play an (in some cases, more) important role at a local scale.

To the east, both Spalding et al (2007) and Obura (2012) placed a latitudinal ecoregion boundary between the Mascarene Islands (Mauritius, Rodrigues and Réunion) in the south; and St Brandon, Tromelin and the Agaléga Islands in the north. Indeed, due to the predominantly zonal currents in this region (the South Equatorial Current), we do not expect strong larval dispersal between these groups of islands. However, due to the seasonal position of the South Equatorial Current and the considerable geographic isolation, our results suggest that the primary latitudinal dispersal barrier is around 15 *^◦^*S (north of St Brandon and Tromelin), rather than further to the south as suggested by Spalding et al (2007) and Obura (2012). Obura (2012) further assigned the Farquhar Group (Seychelles) to the ecoregion containing St Brandon, Tromelin and the Agaléga Islands, but our simulations predict that the Farquhar Group is more strongly connected to the Aldabra Group, as the latter is directly downstream of the former. Burt et al (2024) recently found strong genetic connectivity across Seychelles which, in combination with our simulations, suggests that ocean currents cause Seychelles to act as a distinct biogeographic unit (even for corals with a short pelagic larval duration).

Due to the bifurcation of the South Equatorial Current and the patchiness of coral reefs, we predict a dispersal barrier in northeast Madagascar around 15 *^◦^*S. This is about 3 *^◦^* and 1 *^◦^* north of the analogous ecoregion boundaries identified by Spalding et al (2007) and Obura (2012), respectively. A possible explanation is related to seasonal variability in the bifurcation latitude of the South Equatorial Current: we only consider the northwest monsoon in this analysis (in line with coral spawning seasonality), but bifurcation shifts to the south during the southeast monsoon (Chen et al, 2014), which could affect the dispersal of other taxa, and hence the Spalding et al (2007) ecoregions. However, van der Ven et al (2022) identified strong genetic connectivity for *Acropora tenuis* across the northeast Madagascar barrier identified by our study (as well as the slightly more southerly location suggested by Obura (2012)), instead supporting a barrier more consistent with Spalding et al (2007) (supplementary materials, fig. S13). This could suggest that environmental factors allow strong genetic connectivity to be maintained across northeast Madagascar despite limited dispersal, or that larval parameters (such as spawning seasonality or mortality rate) assumed by our simulations are inappropriate for *Acropora tenuis*. Indeed, this is one of the few dispersal barriers in our simulation that we found to be sensitive to larval parameters, particularly for larvae with lower dispersal potential.

In general, the good agreement between physical dispersal barriers identified by our analysis, biogeographic ecoregions identified by Spalding et al (2007) and Obura (2012), and coral population structure confirms the important role of ocean currents in shaping biodiversity and genetic connectivity. Nevertheless, our findings also highlight that some biogeographic barriers may be maintained despite the potential for strong gene flow (e.g. between the Farquhar Group and the rest of Seychelles), and that physical dispersal barriers do not consistently translate into realised differentiation (e.g. the bifurcation of the Northeast Madagascar Current).

### 4.3 Limitations

Despite the high resolution of our simulations compared to previous studies in the region, we are still, in practice, limited to inferring connectivity at scales of tens of kilometres and above. We therefore do not resolve connectivity (including vertical connectivity) within reef systems (e.g. Saint-Amand et al, 2023; Takeyasu et al, 2023). In this study, most structure in the coral reef networks emerging from multi-generational larval dispersal is at scales of 100 km or greater. This may suggest that the resolution of SECoW is sufficient to reasonably predict these networks, particularly since predicting finer-scale structure may be fundamentally limited due to stochastic variability in ocean currents (Vogt-Vincent et al, 2023). This does, however, raise the question of whether a lower resolution model would be equally suited for predicting large-scale biogeographic patterns. Although raising the model resolution from 1/12*^◦^* (the resolution of many existing global ocean reanalysis) to the 1/50*^◦^* resolution of the present study significantly affects marine dispersal (Poje et al, 2010; Lévy et al, 2012), it is not clear how important this is for connectivity over many generations.

On the other hand, the inability of our model to reproduce flow around coral reefs could conceivably affect the resulting large-scale network structure. Dauhajre et al (2019) showed that models with a comparable resolution to ours underestimate alongshore transport, so our simulations may underestimate connectivity along continental coastlines. Drifter trajectories suggest that a failure to resolve reef-scale oceanographic processes likely overestimates cross-shore transport (and therefore connectivity) between coral islands (Monismith et al, 2018). Reef-scale connectivity simulations have largely been limited to 2D hydrodynamic models investigating dispersal in shallow water (Grimaldi et al, 2022; Saint-Amand et al, 2023), in contrast to the 3D hydrodynamic model used in this study, necessitated by the much larger spatial scale, and larval transport across deep ocean. Nevertheless, these studies suggest that SECoW may overestimate the connectivity of remote reefs, which may have the consequence of increasing the connectivity timescale between reefs (supplementary materials, fig. S14). In a recent study, however, SECoW *underestimated* the realised coral connectivity between remote reefs in Seychelles, as inferred from population genetics (Burt et al, 2024). Regardless, although the inability of the model to reproduce reef-scale dynamics may be of lesser consequence for the large-scale biogeographic patterns described in this study, it is undoubtedly an important consideration at smaller scales. Whilst our results may be useful for marine spatial planning and other conservation applications, we remind readers that they are only valid at spatial scales much greater than the model resolution.

Our model predicts that *Platygyra daedalea* populations in the southwest Indian Ocean are connected over a timescale of around 200 generations (or roughly 1000 years), and longer for corals with a shorter pelagic larval duration. Our multigenerational connectivity networks therefore assume that recent surface currents (as used in this study) are representative of surface currents throughout the late, and potentially mid, Holocene. Based on the southward contraction of negative wind stress curl over the southwest Indian Ocean during the mid-Holocene predicted by PMIP4 models (Guo et al (2019); Mauritsen et al (2019); Hajima et al (2020); Zheng et al (2020); and supplementary materials, fig. S15), it is possible that the South Equatorial Countercurrent previously separated from East Africa at a more southerly latitude during the spawning season. However, given the generally muted response of monsoons in the southwest Indian Ocean to mid-Holocene climate change (Brierley et al, 2020; Leupold et al, 2023), we consider it unlikely that these changes would have substantially changed connectivity networks. Whilst our predictions of coral reef connectivity may therefore be robust with respect to mid-to-late Holocene climate change, western Indian Ocean biogeography has its roots in the Palaeogene, where the tectonic configuration - and therefore surface currents - were considerably different (e.g. Obura, 2016). More recently, much of the Mascarene Plateau may have been exposed during the last glacial maximum, around 21,000 years ago (supplementary materials, fig. S16; Cacciapaglia et al (2021)). If surrounded by fringing reefs, for instance, we could hypothesise that the glacial Mascarene Plateau acted as a stepping stone to facilitate larval dispersal from the Chagos Archipelago to the Mozambique Channel. Patterns of connectivity will therefore undoubtedly have changed throughout geological time, likely explaining some of the differences in biogeography and population structure, and our model predictions.

Finally, although our predictions are generally insensitive to larval competency and mortality parameters, other biological assumptions (for example, that larvae maintain their vertical position near the surface; that there is no significant vertical connectivity with mesophotic reefs that are not included in SECoW; and that fecundity, habitat suitability, and post-settlement mortality are spatially invariant) could affect the network structure. The relative importance of these biological and physical uncertainties is not yet clear, and would benefit enormously from future connectivity studies combining data from numerical models and observations at coherent spatial and temporal scales (Edmunds et al, 2018).

## 5 Conclusions

Despite the presence of many remote islands, our findings suggest that coral reefs across the tropical southwest Indian Ocean are strongly connected over evolutionary timescales, with most reefs physically connected within a few hundred generations.

Our larval dispersal simulations suggest that the Northeast Madagascar Current acts as a leaky dispersal barrier, separating Seychelles in the north from the Mozambique Channel in the south, before bifurcating at the East African coast and generating a further barrier to larval dispersal between Mozambique and Tanzania. The East African Coastal Current facilitates strong and consistent connectivity along the coasts of Tanzania and Kenya, with a secondary dispersal barrier generated in the wake of Mafia Island. The Chagos Archipelago, whilst geographically isolated, is in a position to receive significant numbers of larvae from the Inner Islands of Seychelles through the ephemeral surface expression of the Southeast Equatorial Countercurrent, and broadcasts larvae to the Mozambique Channel through the South Equatorial Current. Although the flow of larvae between distant populations is generally unlikely to be high enough to drive demographic change, it is certainly strong enough to establish long-distance genetic connectivity (e.g. Lowe and Allendorf, 2010), as confirmed by the genetic and biogeographic studies discussed in this paper. Our findings suggest that the large-scale coral population structure of the tropical southwest Indian Ocean can be explained reasonably well through physical larval dispersal, with contemporary ocean currents playing a significant role in setting biogeographic barriers and gene flow. However, passive larval dispersal alone is unsurprisingly insufficient to fully explain regional biogeography and population genetics. It is not yet clear to what extent these differences have emerged due to palaeogeography and/or palaeoceanography, complex larval behaviour, local adaptation and other post-settlement processes, or sampling limitations by genetic and biogeographic studies.

To our knowledge, no genetic studies have investigated the relative connectivity of corals between Seychelles, the Chagos Archipelago, the Mascarene Plateau and Islands, and the Mozambique Channel, so there are limited genetic data to contextualise our predictions of connectivity across the many remote islands of the southwest Indian Ocean. A large scale, methodologically consistent population genetics study across the region would first be needed, in order to carry out a quantitative comparison to more rigorously test our model predictions.

Nevertheless, this study represents an important step towards quantifying the connectivity of coral reefs across the southwest Indian Ocean, and we hope that our connectivity predictions, which are freely available in full (Vogt-Vincent et al, 2024), will be useful to those studying biogeography and population structure in the region, by providing quantitative physical context to their observations. Marine managers and governments looking to enhance national or regional coral reef system resilience could also integrate these predictions (and the associated web app) into marine spatial planning efforts, to identify important source and sink reefs, and thereby prioritise conservation efforts.

## 6 Data availability

The connectivity matrices used in this study, as well as all figures and scripts required to reproduce figures from the main text and supplement, are archived and documented in a Zenodo repository (Vogt-Vincent et al, 2024). All hydrodynamic data used in this study are described by Vogt-Vincent and Johnson (2023) and are available at the CEDA Archive. SECoW was described by Vogt-Vincent et al (2023), with the code available in the associated datasets.

## 7 Acknowledgements

This work was funded by Natural Environment Research Council grant NE/S007474/1 and used the ARCHER2 UK National Supercomputing Service (https://www.archer2.ac.uk), and JASMIN, the UK collaborative data analysis facility. We thank the two reviewers for their thoughtful comments and suggestions, which have significantly improved the clarity of the manuscript, and the robustness of our conclusions. We are also grateful to Otis Brunner, who helped conceptualise and develop the Sea the Connectivity web app (used here to interactively visualise the explicit connectivity matrix) with Noam Vogt-Vincent; Julia Janicki, who wrote the underlying code for Sea the Connectivity; and the SDG Fund at the Okinawa Institute of Science and Technology, which funded the development of this app.

## 8 Conflict of interest statement

On behalf of all authors, the corresponding author states that there is no conflict of interest.

## Notes

https://zenodo.org/doi/10.5281/zenodo.10183948

### Competing Interest Statement

The authors have declared no competing interest.

### Summary of Updates

Manuscript was revised after a first round of peer-review.

