## Supplementary materials for "Coral reef potential connectivity in the southwest Indian Ocean"

### Supplementary Material for: Coral reef potential connectivity in the southwest Indian Ocean

#### 1 Larval dispersal model (SECoW)

A full description of the SECoW larval dispersal model can be found in the supplementary materials of Vogt-Vincent et al (2023), but we provide an overview here for reference, abridged from Vogt-Vincent et al (2023).

SECoW assumes that coral larvae physically behave as positively buoyant, otherwise passive particles, which are advected by surface currents for 120 days after spawning, using a fourth-order explicit Runge-Kutta scheme in OceanParcels (using a time-step of 10 minutes). The ocean current data from WINDS is defined on a C-grid, so particles do not enter land cells aside from rare integration errors. We did not use a parameterisation for sub-grid scale processes, as it is not clear that a simple random walk parameterisation would result in any meaningful improvement at the relatively high resolution and short connectivity timescales used in this study (Okubo, 1971; van Sebille et al, 2018; Reijnders et al, 2022). Using a stochastic parameterisation would also require an additional (and arbitrary) parameterisation to prevent particles from entering land cells, and any uncertainty introduced by not resolving sub-grid scale processes is likely small compared to the biological uncertainties (such as a poor understanding of the vertical positioning of larvae in the open ocean due to active swimming and buoyancy versus passive advection).

We released particles from every c. 2×2 km reef cell on the WINDS grid, based on an upscaled reef cover map derived from satellite imagery (Li et al, 2020), resulting in 8088 reef cells across the WINDS domain. We released $N = 1024$ (32×32) particles per reef cell, arranged in a regular grid, with the particle density derived from a sensitivity analysis (Figure S1, Vogt-Vincent et al (2023)). We then simulated daily spawning events at midnight from October to January (1993 – 2019), for a total of 4920 domain-wide spawning events (a subset of the simulations carried out in Vogt-Vincent et al (2023)), with c. $8.3\times10^6$ particles seeded per event. Each particle represents many coral larvae, assumed to have an initial separation small enough such that they do not disperse significantly relative to one another. In the present study, we assumed a constant coral fecundity per unit area per day, $\rho$. The computation of larval fluxes between cells necessitates a value for $\rho$, which we set to $\rho = 1$ larva m$^{-2}$ d$^{-1}$, but this value does not affect any patterns or trends in this study as long as it is constant. In reality, coral fecundity is certainly not constant (e.g. Hartmann et al, 2018), but this is a necessary approximation for now, as no maps of coral fecundity exist spanning the southwest Indian Ocean. Therefore, each particle released in cell $i$ represents $L_i^0 = \rho A_i/N$ larvae at spawning, where $A_i$ is the reef cover of cell $i$ for the purpose of this study.

Following Connolly and Baird (2010), we assumed that the coral larvae represented by each particle are split across three larval reservoirs: a pre-competent reservoir $L_1$, a competent reservoir $L_2$, and a post-competent or dead larval reservoir $L_3$. The pre-competent and competent reservoirs evolve through time according to the following differential equations:

$$\frac{dL_1}{dt} = -\alpha^*(t)L_1(t) - \mu_m(t)L_1(t)$$
$$\frac{dL_2}{dt} = \alpha^*(t)L_1(t) - \beta L_2(t) - \mu_m(t)L_2(t) - \mu_s F_r(t)L_2(t)$$

$$\alpha^*(t) = \begin{cases} 0 & t < t_c \\ \alpha & t \geq t_c \end{cases}$$

Where $\alpha^*$ is a competency acquisition rate ($\mathrm{d}^{-1}$), $\beta$ is a competency loss rate ($\mathrm{d}^{-1}$), $\mu_m$ is a mortality rate ($\mathrm{d}^{-1}$), $\mu_s$ is a settling rate ($\mathrm{d}^{-1}$), and $F_r$ is the fraction of a WINDS cell area covered by reef. The parameter $a^*$ is equal to 0 before a larva has reached its minimum competency period $t_c$ (d), and $\alpha$ thereafter. The settling term $\mu_s F_r(t)L_2(t)$ was not present in the original formulation of Connolly and Baird (2010). Since SECoW cannot provide any physical insight into the position of a particle below the WINDS grid resolution (c. 2 km), we assumed that the particle is equally likely to be anywhere in the reef cell it currently occupies for the purpose of this biological parameterisation. The settling rate is therefore equal to a fixed settling rate $\mu_s$ assuming 100% reef coverage, multiplied by the actual proportion of the reef cell covered by reef. In this study, we set $\mu_s = 1\ \mathrm{d}^{-1}$, assuming settling times on the order of a day (Tay et al, 2011), although this number is highly uncertain and will in reality depend on reef structural complexity and the availability of settlement cues (Harrington et al, 2004; Hata et al, 2017). Nevertheless, the settling rate parameter primarily affects the absolute number of larvae settling, rather than the distribution of larvae between reefs, and therefore only has a limited impact on network structure (supplementary fig. 17). In contrast to assumptions of instantaneous settling (e.g. Holstein et al, 2016; Figueiredo et al, 2022) or use of a Lagrangian Probability Density Function (LPDF, see Mitarai et al (2009) and applications in Uchiyama et al (2018); Thompson et al (2018)), this method accounts for the capacity of upstream reefs to reduce the larval supply to downstream reefs (contrary to the LPDF), whilst sensibly handling sub-grid scale reef coverage and allowing for the possibility of a larva passing over a reef without settling (Hata et al, 2017). We solve for the number of larvae $N_s$ settling during each settling event $i$ by solving $N_s^i = \mu_s F_r^i \int_t^{t+\Delta t} L_2\ dt$, where $t$ and $\Delta t$ are respectively the start time and duration of the settling event.

#### 2 Influence of mortality and competency dynamics

The results discussed in the main text use larval parameters based on *Platygyra daedalea*, but we have repeated the analyses based on four other coral species (Connolly and Baird, 2010) (see Supplementary Fig. 18 and Supplementary Table 1 for further details). The raw data and figures for all five species are available in the supplementary dataset. Here, we briefly discuss the similarities and differences.

| Species | Code | $\alpha^*$ ($d^{-1}$) | $\beta$ ($d^{-1}$) | $t_c$ (d) | $\lambda$ ($d^{-1}$) | $\nu$ | $\sigma$ | $\mu_s$ ($d^{-1}$) |
|---|---|---|---|---|---|---|---|---|
| *A millepora* | AM | 0.18 | 0.050 | 3.239 | 0.043 | 0.57 | 0.0 | 1.0* |
| *A valida* | AV | 0.22 | 0.031 | 1.0* | 0.019 | 0.46 | 0.0 | 1.0* |
| *A gemmifera* | AG | 0.39 | 0.145 | 3.471 | 0.067 | 1.0 | 0.0 | 1.0* |
| *G retiformis* | GR | 0.58 | 0.096 | 1.0* | 0.087 | 1.0 | 0.0 | 1.0* |
| *P daedalea* | PD | 0.39 | 0.099 | 2.937 | 0.060 | 0.72 | 0.0 | 1.0* |
| *P daedalea (rapid settling)* | PD-6S | 0.39 | 0.099 | 2.937 | 0.060 | 0.72 | 0.0 | 6.0* |

**Supplementary Table 1** Biological parameters used for larval competency, mortality and settling in this study: $\alpha$ (competency acquisition rate), $\beta$ (competency loss rate), $t_c$ (minimum competency period), mortality parameters $\lambda$, $\nu$ and $\sigma$ (see Connolly and Baird (2010)), and settling rate $\mu_s$. An asterisk indicates that the parameter used is not identical to Connolly and Baird (2010). For the minimum competency period $t_c$, this is due to the minimum of 1 day imposed by SECoW. For the settling rate $\mu_s$, this is because settling rate was not included in the model of Connolly and Baird (2010). *Platygyra daedalea* is used for most results in this study. This table is adapted from Table S1 in Vogt-Vincent et al (2023).

The single-step explicit and implicit connectivity matrices are qualitatively similar across species, although the absolute values of explicit connectivity are predictably higher for the species with the lowest larval mortality rates (with the difference being most extreme for sites separated by the greatest oceanographic distances). For instance, the likelihood of a *Goniastrea retiformis* larva from a reef in the Chagos Archipelago reaching Madagascar is less than $10^{-9}$, whereas the same probability for *Acropora valida* is up to $10^{-5}$.

Similarly, fewer steps of dispersal tend to separate distant pairs of reefs through the implicit connectivity metric, for larvae with lower mortality rates and a longer competency period. Aside from the Mascarene islands, we expect that almost all reefs in the southwest Indian Ocean physically share an ancestral reef within around 200 steps of dispersal for *Platygyra daedalea* (see main text). This falls to around 100 steps of dispersal for long-lived *Acropora valida* larvae, and over 500 for *Goniastrea retiformis* (particularly for the Chagos Archipelago). Implicit connectivity between nearby sites is often higher for coral species with shorter pelagic larval duration. This is because the diversity of larval destinations is lower, with gene flow tending to be retained within small-scale local reef clusters (rather than 'leaking' to distant reefs).

There are some minor differences in the diversity of larval destinations (*out-entropy*, see Supplementary Fig. 4) across the five species. Unsurprisingly, larvae with a longer competency window tend to reach a greater diversity of destination reefs, a pattern that is most pronounced along the path of the East African Coastal Current and

Northeast Madagascar Current. Interestingly, *Goniastrea retiformis* larvae from the Chagos Archipelago reach a high diversity of reefs, despite having one of the shortest competency windows amongst the five tested species. This is likely because *Goniastrea retiformis* larvae reach peak competency the fastest, and due to the remoteness of the Chagos Archipelago, almost all realistic larval destinations are within the archipelago. A longer competency window is therefore of limited use if the larvae do not mature quickly enough to settle within the archipelago before they are swept out to the open ocean. A short larval duration may therefore be beneficial in this case. Longer competency windows and lower mortality rates also (again, unsurprisingly) increase the overall proportion of larvae that settle anywhere, again with the exception of *Goniastrea retiformis* at some remote reefs. T

The dominant principal component (PC1) in the meta-clustering analysis is practically identical across the five species, representing the consistent and strong connectivity along the coast of East Africa facilitated by the East African Coastal Current. PC2 is broadly similar, representing connectivity between the Mozambique Channel region and the Chagos Archipelago, but the Chagos Archipelago has higher values for coral species with longer competency windows, representing more consistent connectivity. PC3 shows two different patterns. For larvae with intermediate larval duration (*Platygyra daedalea*, *Acropora gemmifera*, and *Acropora millepora*), PC3 is as shown in Supplementary Fig. 11, representing connectivity between the South Equatorial Current and the East African Coastal Current. For the larvae with the shortest and longest larval duration (*Goniastrea retiformis* and *Acropora valida*), PC3 instead appears to represent the role of the South Equatorial Current as a dispersal barrier. It is not clear whether this difference is meaningful, or whether this just represents a small difference between lower-order principal components, exacerbated by the decision to focus on the top three.

By automatically partitioning the domain into 8 meta-clusters using through K-Means clustering, all five species show very similar patterns. All display major multi-generational dispersal barriers between Mozambique and Tanzania; at Mafia Island; across the Northeast Madagascar Current; and across the Southeast Equatorial Current. In contrast to the longer-lived larvae (including *Platygyra daedalea*) where Seychelles is split into two clusters, the entire of Seychelles is grouped into a single cluster for *Acropora gemmifera* and *Goniastrea retiformis*. Instead, for these shorter-lived larvae, complex structure appears within the east Mozambique Channel, indicating that connectivity along the west coast of Madagascar in particular may be considerably reduced for larvae with a shorter pelagic larval duration. This may be due to the fragmented reef habitat, complex coastline, and coastal currents favouring retention during spawning months, potentially disproportionately affecting short-lived larvae. For *Goniastrea retiformis* (the species with the shortest lived larva), a dispersal barrier appears in northern Kenya, at precisely the location of the ecoregion boundary identified by Spalding et al (2007) (Supplementary Fig. 12). This indicates that the difference in biogeography across this boundary for a broader range of taxa may be strongly influenced by those with a shorter larval duration. Conversely, *Goniastrea retiformis* appears less affected by the dispersal barrier introduced by the bifurcation of the Northeast and Southeast Madagascar Current in eastern Madagascar. However, this is probably due to a relative increase in the importance of other dispersal barriers, rather than an actual increase in larvae crossing this eastern Madagascan dispersal barrier.

### 3 Infomap and the Map Equation

Consider a random walk through the coral reef network, where the probability of the walk transiting from node $i$ to node $j$ is proportional to the explicit, single-step connectivity between them. A simple approach to describing this random walk would involve assigning a unique code to each node, and representing the random walk as a list of codes. We have 180 reef nodes in our network, so $\lceil \log_2 180 \rceil = 8$ bits of information are required to describe every node visited.

This strategy is inefficient, since some nodes are more likely to be visited than other nodes. An alternative approach might involve simulating large numbers of random walks, observing how many times each node is visited, and assigning Huffman codes based on these visitation probabilities. This involves assigning shorter codes to frequently visited nodes, and longer codes to rarely visited nodes, resulting in an overall reduction in the information required to describe a random sequence of nodes (Huffman, 1952). However, this approach is still not optimal for describing a random walk through most realistic networks. This is because many networks are partitioned into clusters that tend to retain a random walk (or flow) for an extended period of time, so the likelihood of the random walk transitioning to node $j$ is strongly dependent on the identity of node $i$. As an example, whilst a reef along the coast of Tanzania may appear many times in an arbitrarily long random walk, a random walker that is currently in the Chagos Archipelago is extremely likely to remain within the Chagos Archipelago across a large number of steps.

Instead, we could partition the network into clusters that tend to retain random walks, and assign Huffman codes to nodes based on visitation frequency within the cluster (Rosvall et al, 2009). Since codes are now no longer unique to individual nodes (only being unique within the scope of a cluster), the number of bits required to describe a node is reduced. To deal with this non-uniqueness, codes are assigned to clusters, and the cluster code is reported when the random walker moves between clusters. However, as long as these clusters represent collections of nodes that tend to retain random walks, random walks will only rarely move between clusters, and the information required to describe a random walk is reduced (due to the reduction in information required to describe a random walk within clusters). The more clusters there are, the fewer nodes there are per cluster (and therefore the shorter node codes are). However, more clusters also increase the length of cluster codes, and the frequency at which they have to be used. The map equation (Rosvall et al, 2009) and Infomap algorithm (Edler et al, 2023) identifies this optimal partitioning of the network. Note that partitioning can also be hierarchical (i.e. partitioning clusters into sub-clusters, and so on).

Since we can model the movement of genes as a random walk through the network (for now neglecting complications such as effects of selection), this optimisation problem identifies clusters of reefs that would be expected to retain larvae and gene flow over a significant number of generations and spawning events, based on potential connectivity.

A widely used algorithm to partition ecological networks is *modularity* (e.g. Gamoyo et al, 2019; Brunner et al, 2022; Treml and Halpin, 2012; Lequeux et al, 2018; Frys et al, 2020; Gabriela Mayorga-Adame et al, 2022), which maximises the total weight of links within a cluster minus the weight of links with nodes outside the cluster (identifying clusters with strong intraconnectivity and weak interconnectivity). Intuitively, this sounds as if this algorithm would be very well suited to identify clusters of reefs that tend to retain larvae. However, consider Supplementary Fig. 19 (Rosvall et al, 2009). Supplementary Fig. 19(a,b) show two possible partitions of a flow network. Through inspection of this flow network, it is obvious that flow will tend to be retained within the cycles in each corner, since leaving a cycle is only possible within a quarter of nodes. We may therefore expect that this network would be partitioned into four clusters, one for each cycle (Supplementary Fig. 19(a)), and indeed this is the optimal clustering identified by both the map equation and modularity, versus one single cluster for the entire network (Supplementary Fig. 19(b)).

For the networks represented by Supplementary Fig. 19(c,d), flow through the network is no longer possible: the longest random walk has length 2. There is no partitioning of the network that retains flow because a random walk will immediately enter a sink node, regardless of where it is within the network. This is recognised by the map equation, but modularity identifies the same partition as in Supplementary Fig. 19(a), since the total weight of links within clusters versus between clusters is unchanged.

### Supplementary Figures

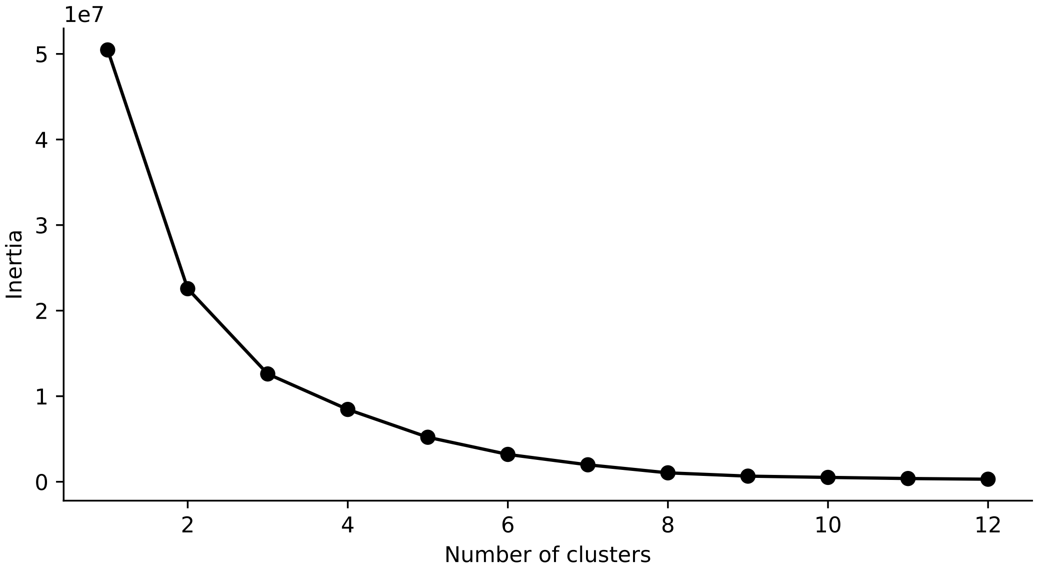


**Supplementary Fig. 1** Inertia (sum of the squared distances (in principal component space) of reefs to their assigned cluster centroid) as a function of the number of K-Means clusters, for *Platygyra daedalea*. Note that the $y$ axis is scaled by a factor of $10^7$.

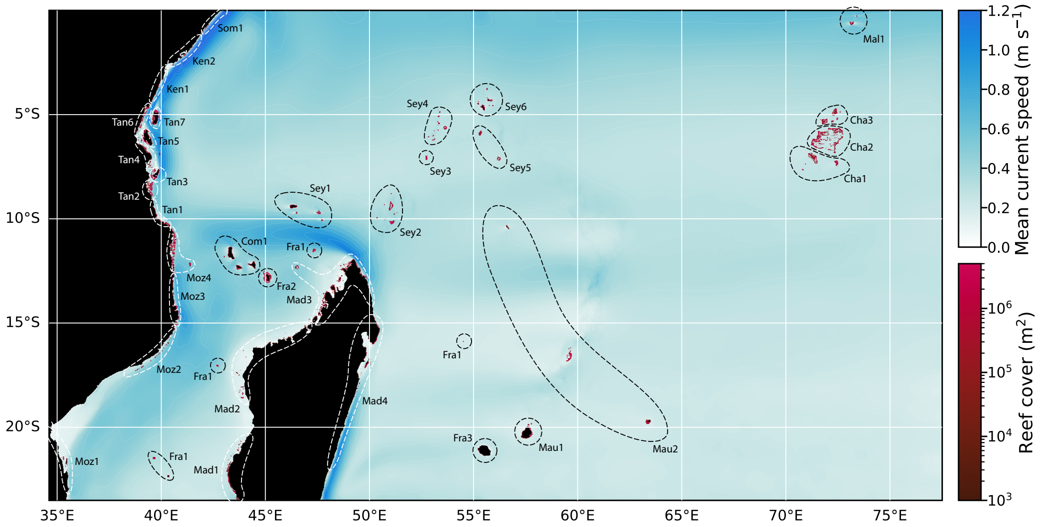


**Supplementary Fig. 2** Key to subregion codes in fig. 2 (main text).

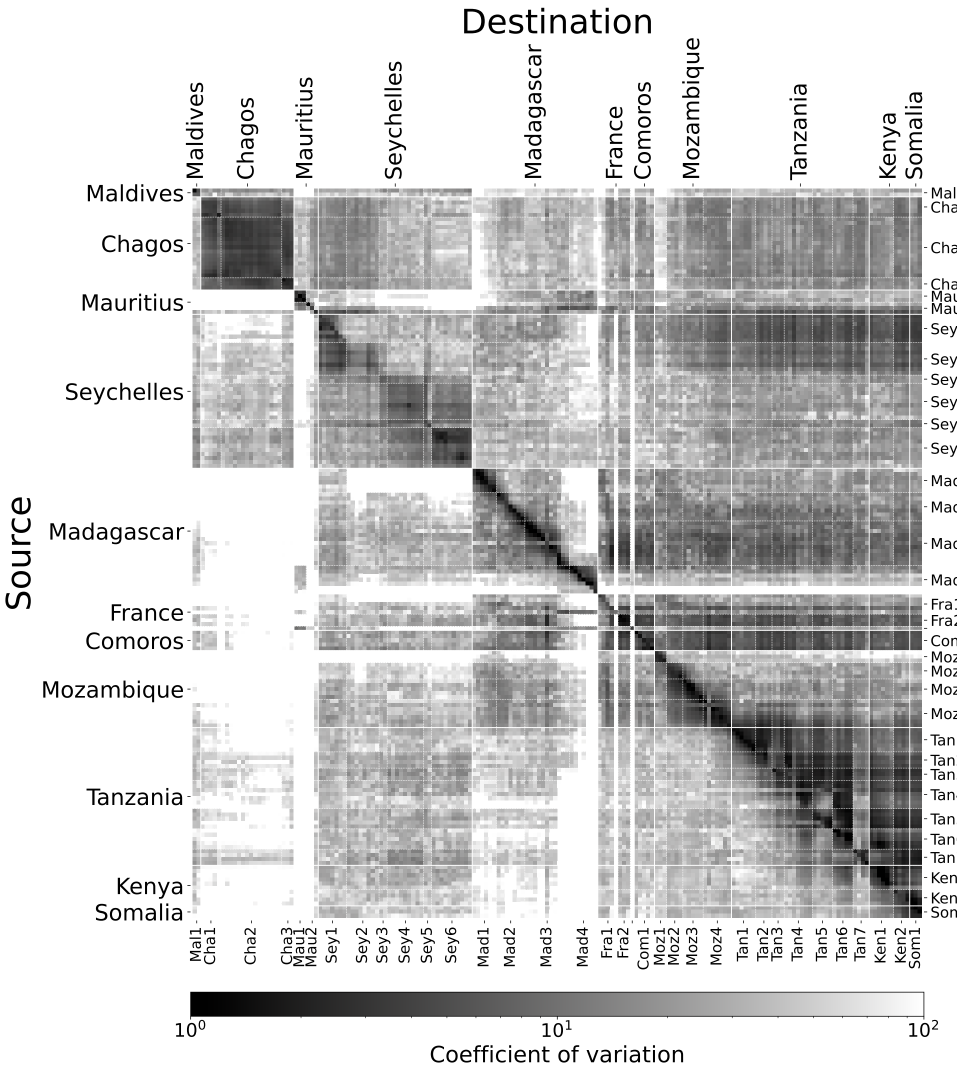


**Supplementary Fig. 3** Coefficient of variation (standard deviation divided by the mean) of potential connectivity between all pairs of reef groups. Subregion codes are as follows: **Mal1**: Maldives; **Cha1**: S Banks; **Cha2**: Grand Chagos Banks; **Cha3**: N Banks; **Mau1**: Mauritius (Island); **Mau2**: Outer Islands; **Sey1**: Aldabra Grp; **Sey2**: Farquhar Grp; **Sey3**: Alphonse Grp; **Sey4**: Amirantes; **Sey5**: Southern Coral Grp; **Sey6**: Inner Islands; **Mad1**: SW Madagascar; **Mad2**: NW Madagascar; **Mad3**: N Madagascar; **Mad4**: E Madagascar; **Fra1**: Scattered Islands; **Fra2**: Mayotte; **Fra3**: Réunion (not labelled); **Com1**: Comoros; **Moz1**: S Mozambique; **Moz2**: Primeiras & Segundas; **Moz3**: N Mozambique; **Moz4**: Quirimbas; **Tan1**: S Tanzania; **Tan2**: Songo Songo; **Tan3**: Mafia Island; **Tan4**: Dar es Salaam region; **Tan5**: Zanzibar; **Tan6**: Tanga; **Tan7**: Pemba; **Ken1**: S Kenya; **Ken2**: N Kenya; **Som1**: S Somalia

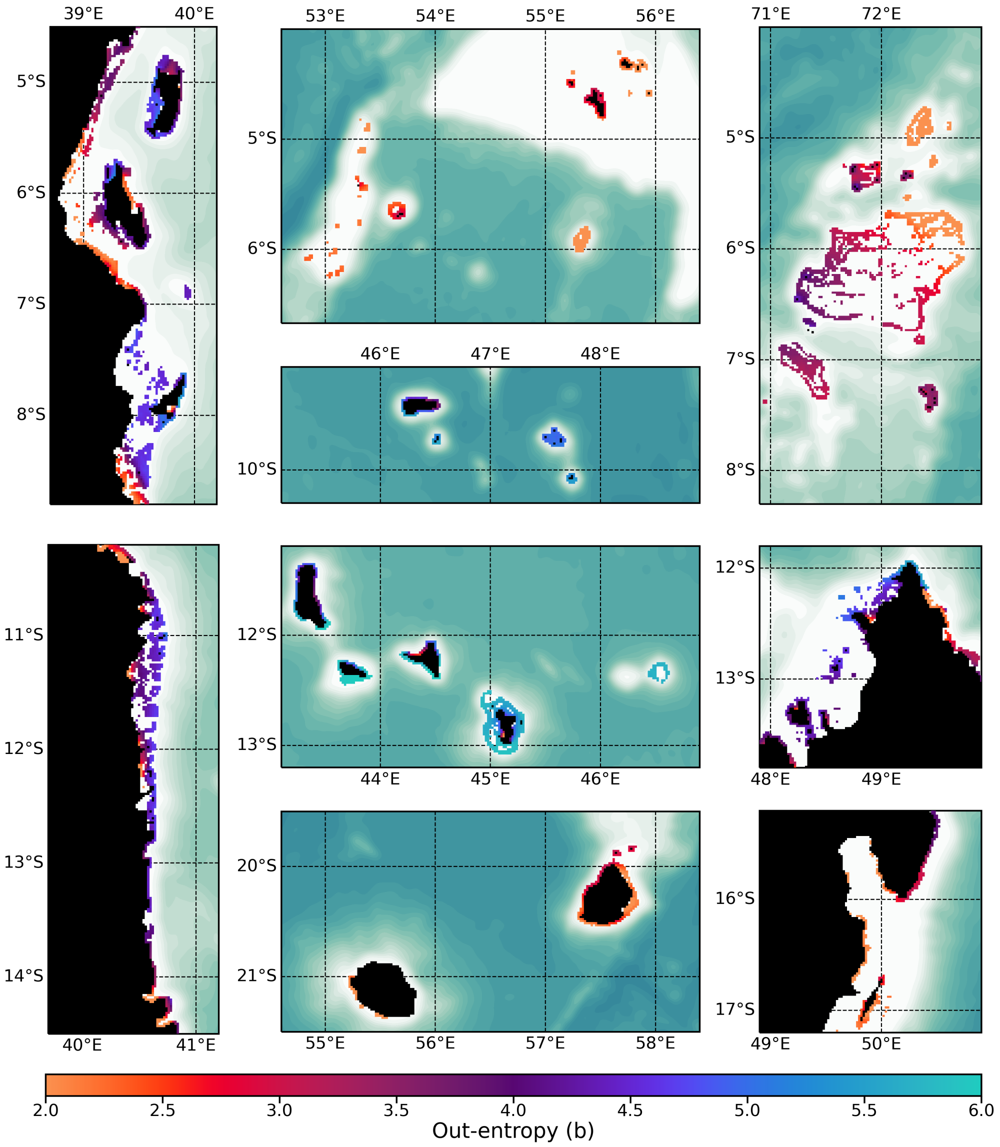


**Supplementary Fig. 4** Median *out-entropy* across daily potential connectivity matrices. We define the out-entropy $H_{ik}$ as the entropy associated with larval destinations $j$ for larval source $i$ on day $k$, $H_{ik} = -\sum_j \left( C'_{ijk} \log_2 C'_{ijk} \right)$, where $C'_{ijk} = C_{ijk} / \sum_j C_{ijk}$. Out-entropy is maximised by larvae being distributed in similar numbers across many destination reefs. An out-entropy of $h$ can be interpreted as the equivalent of a spawning event distributing larvae in equal numbers across $2^h$ reefs. Shown here (clockwise from top-left) is the coast of Tanzania (with Mafia, Zanzibar, and Pemba islands); the Inner Islands of Seychelles and Amirantes; the Aldabra Group; the Chagos Archipelago; north Madagascar; east Madagascar; Réunion and Mauritius; the Comoro Islands; and north Mozambique and the Quirimbas Archipelago.

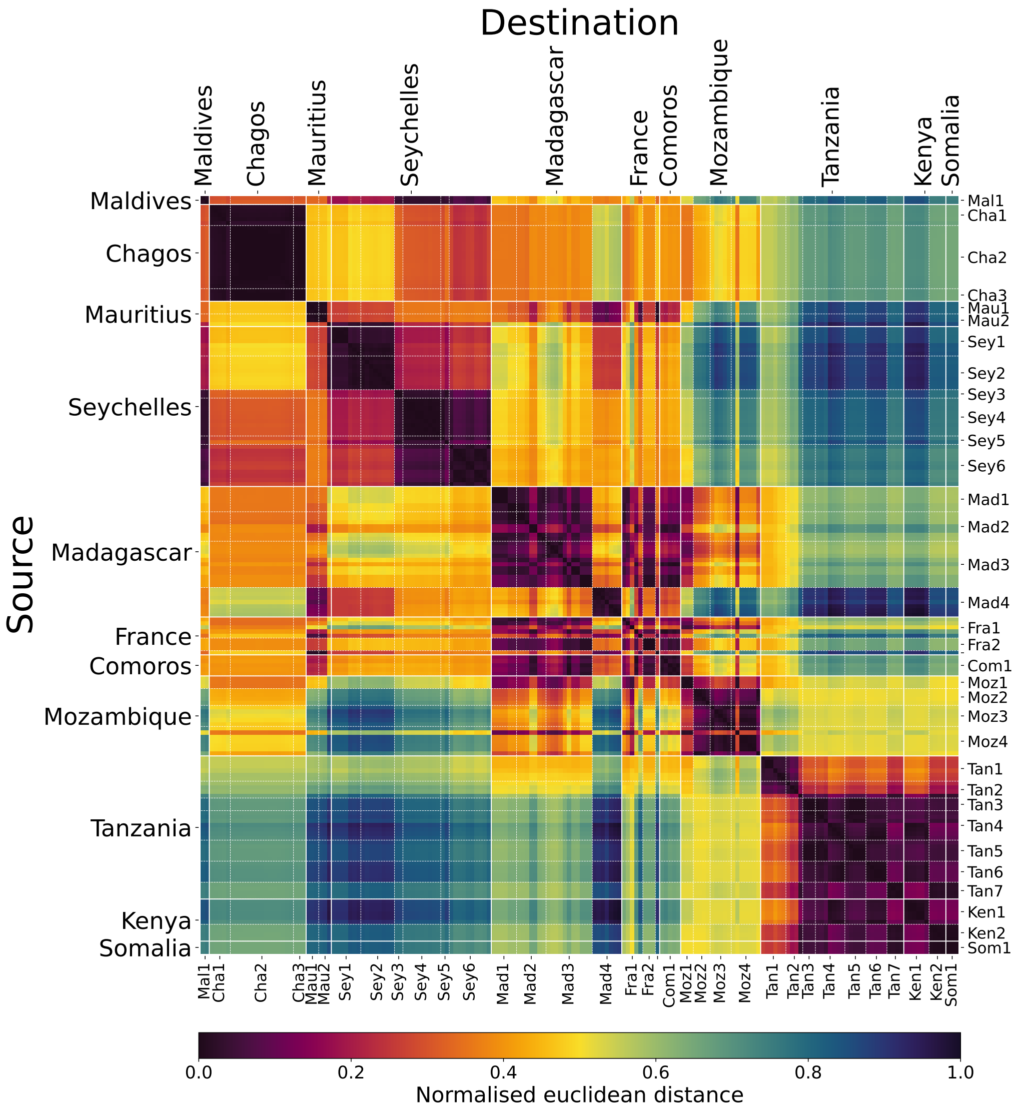


**Supplementary Fig. 5** Euclidean distance between coral reef groups embedded in PC1-PC2-PC3 space (scaled between 0 and 1), where low distance indicates that reef groups are consistently assigned to similar clusters by the Infomap algorithm across stochastic oceanographic variability (based on dispersal across $k = 10$ dispersal events, as in the main text). Note that principal components are normalised for clarity to zero mean and unity standard deviation in the main text, but the distances in this figure are based on the true Euclidean distance. Subregion codes as in Supplementary Fig. 3.

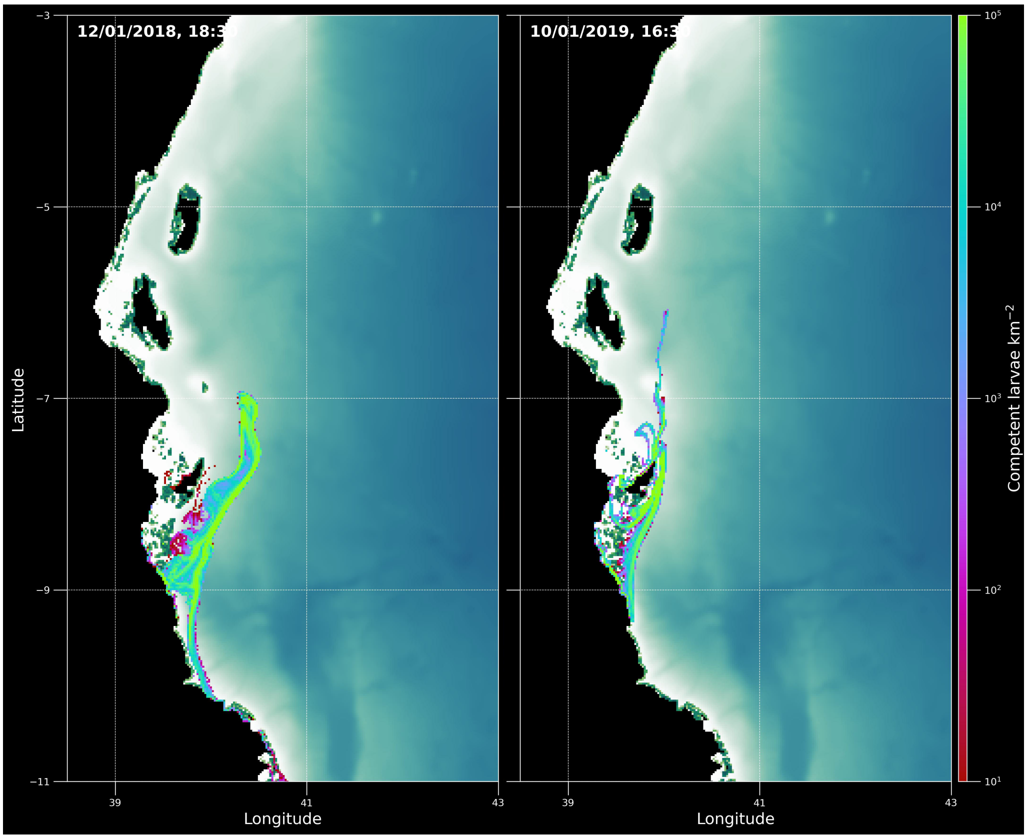


**Supplementary Fig. 6** Two snapshots of coral larval density (based on a scaled-down version of our full larval dispersal simulations, run for visualisation purposes) for simulated spawning events upstream of Mafia Island on 1[st] January 2018 and 2019.

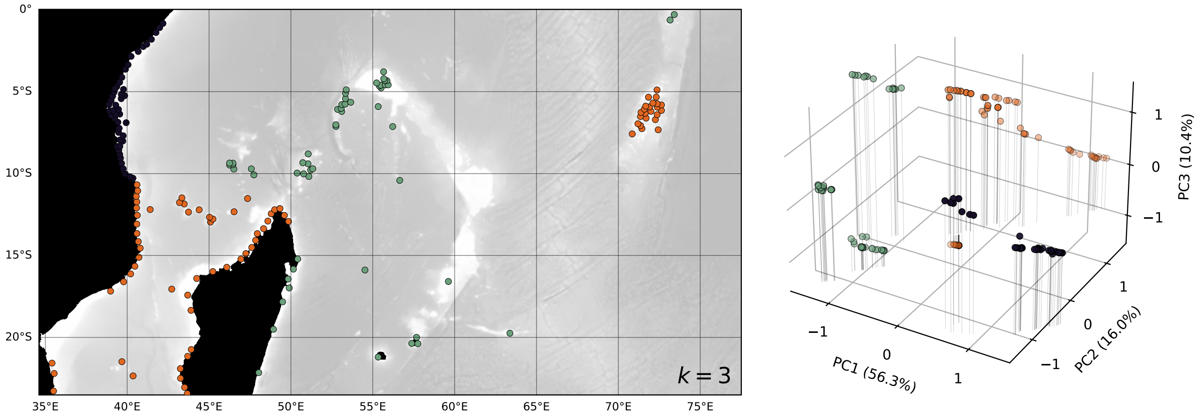


**Supplementary Fig. 7** Reefs plotted geographically and in PC space (as in fig. 4-5 in the main text) but coloured discretely based on cluster assignment by the K-Means algorithm, using 3 clusters.

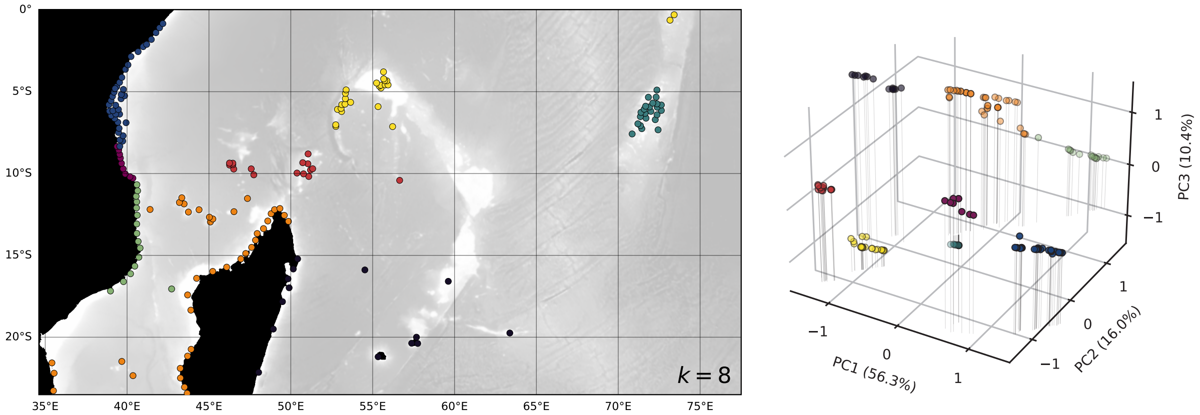


**Supplementary Fig. 8** As in supplementary fig. 7, but for 8 clusters.

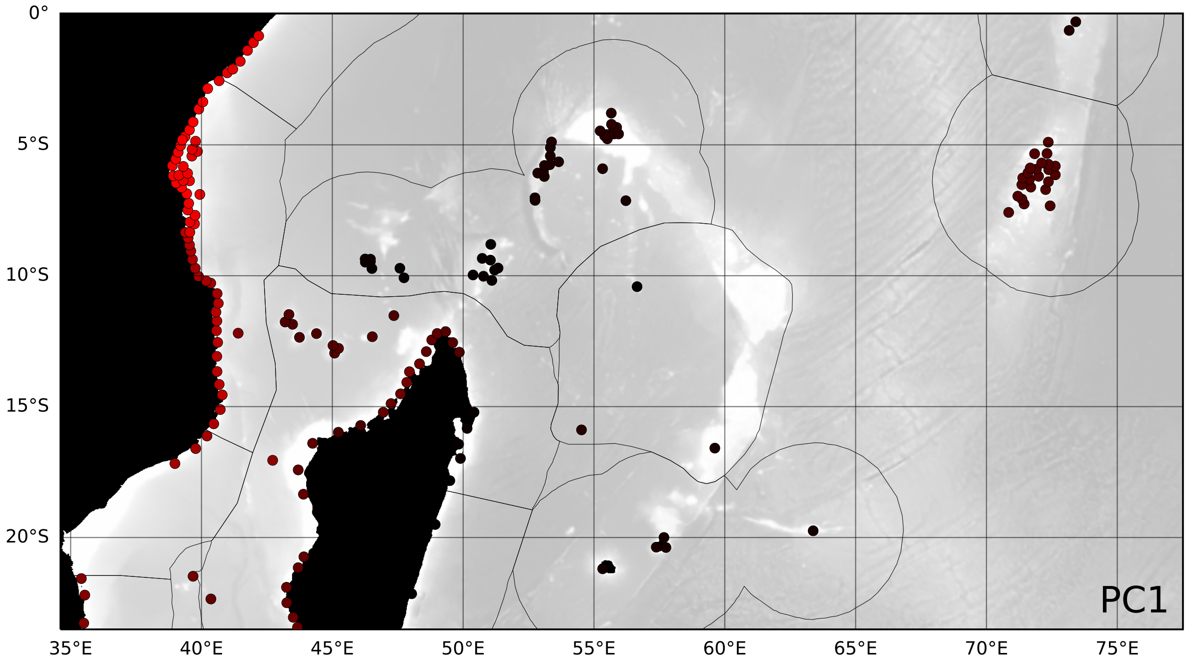


**Supplementary Fig. 9** Reefs coloured by PC1 only.

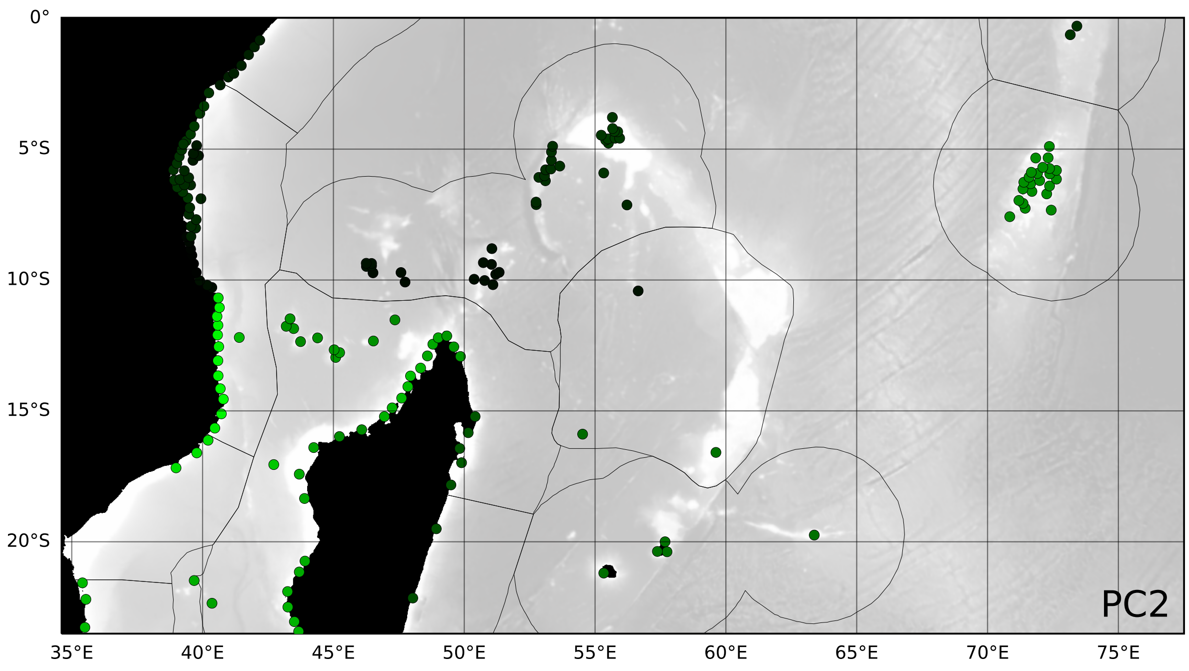


**Supplementary Fig. 10** Reefs coloured by PC2 only.

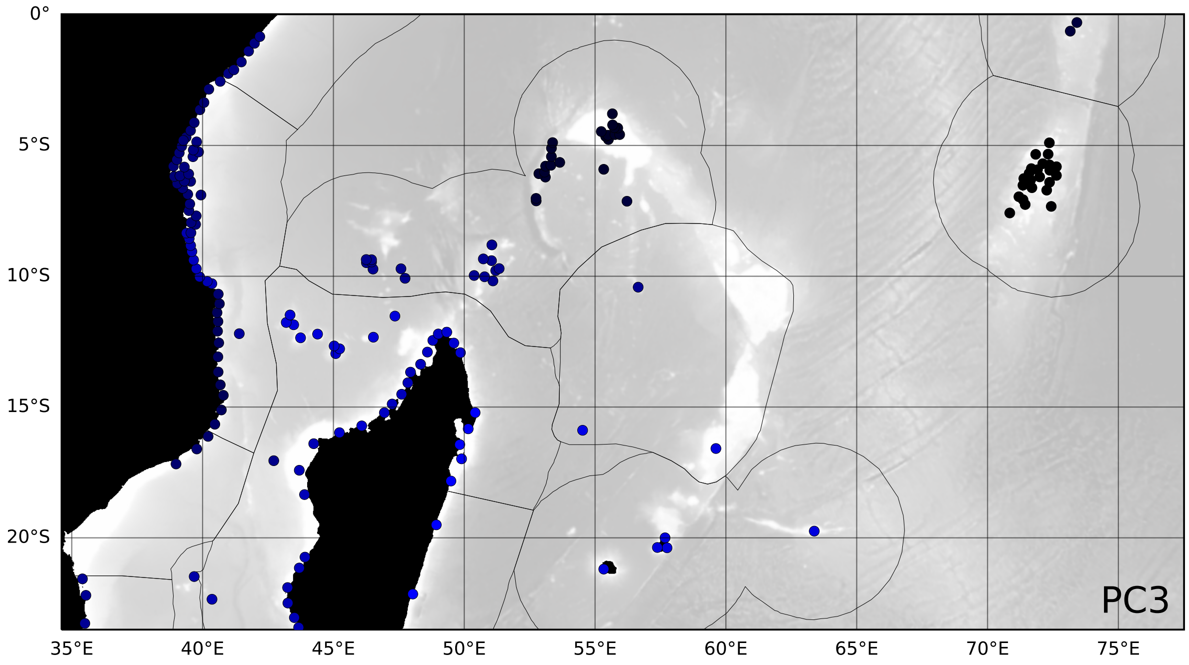


**Supplementary Fig. 11** Reefs coloured by PC3 only.

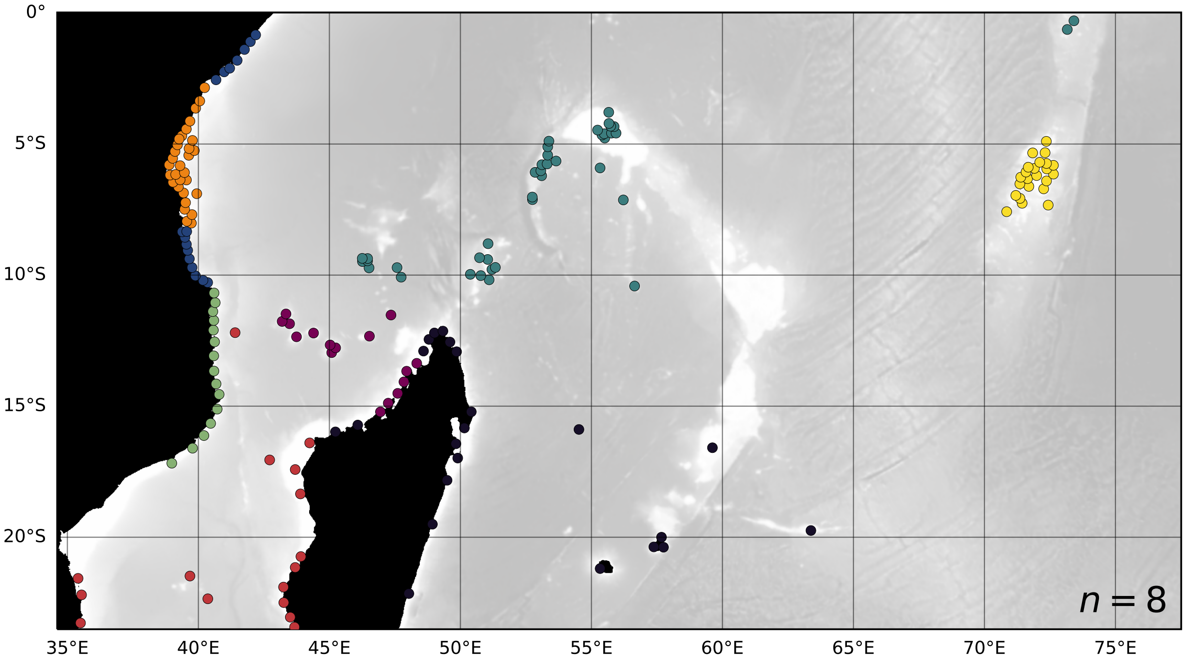


**Supplementary Fig. 12** As in supplementary fig. 8, but for *Goniastrea retiformis*.

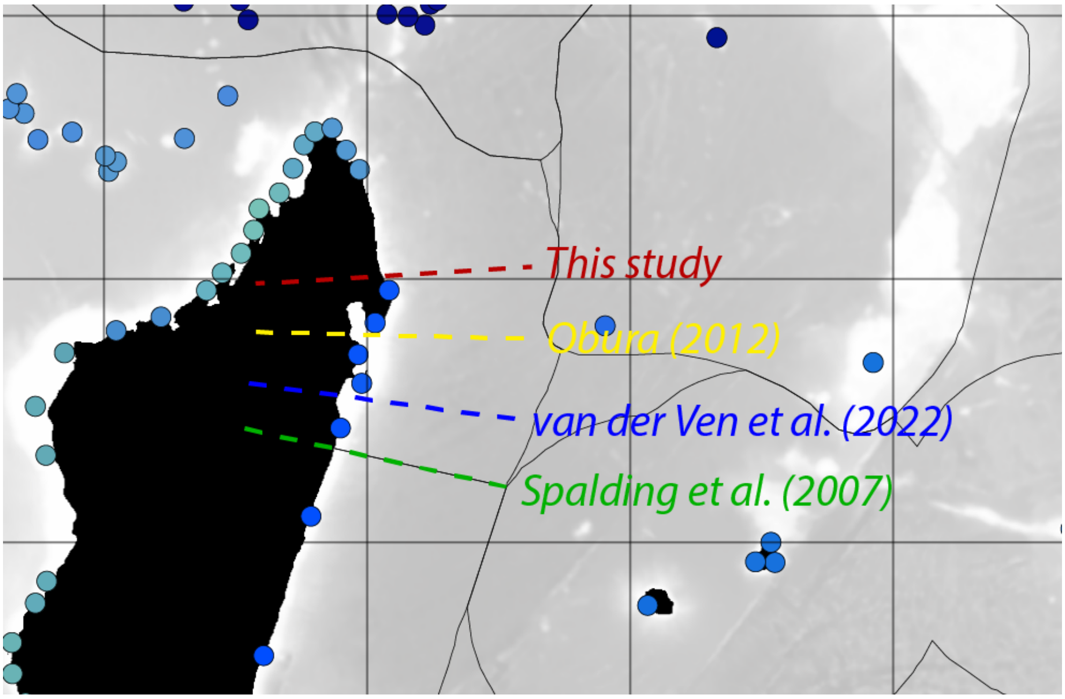


**Supplementary Fig. 13** Approximate locations of the northeast Madagascar dispersal/biogeographic barrier identified by this study (red), Obura (2012) (yellow), van der Ven et al (2022) (blue), and Spalding et al (2007) (green). Note that the barrier locations indicated in this diagram from this study and van der Ven et al (2022) are respectively southernmost and northernmost bounds, rather than exact locations.

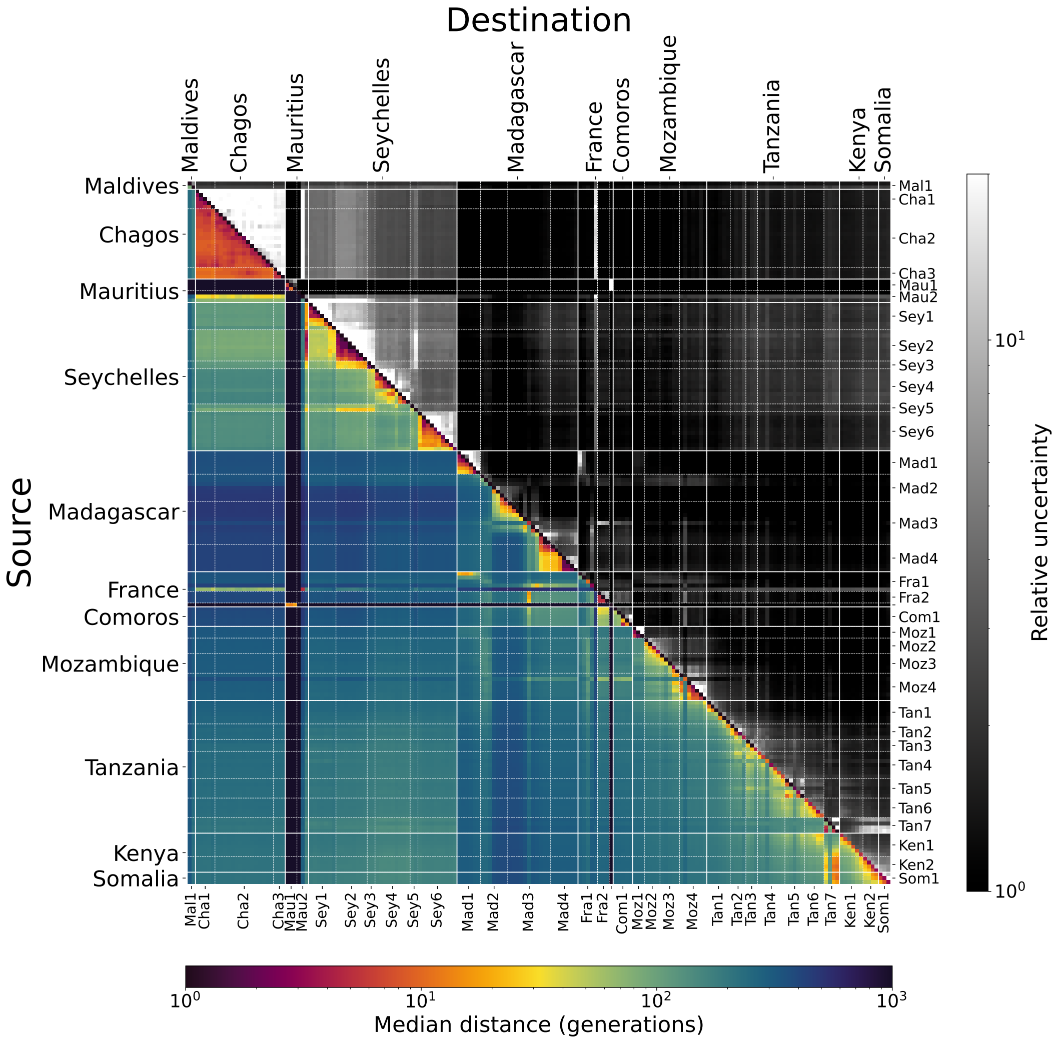


**Supplementary Fig. 14** As in Fig. 3 in the main text, but decreasing the proportion of larvae connected to other reef groups by a factor of 10 (i.e. increasing relative retention by a factor of 10). Note the different scale for the colour map.

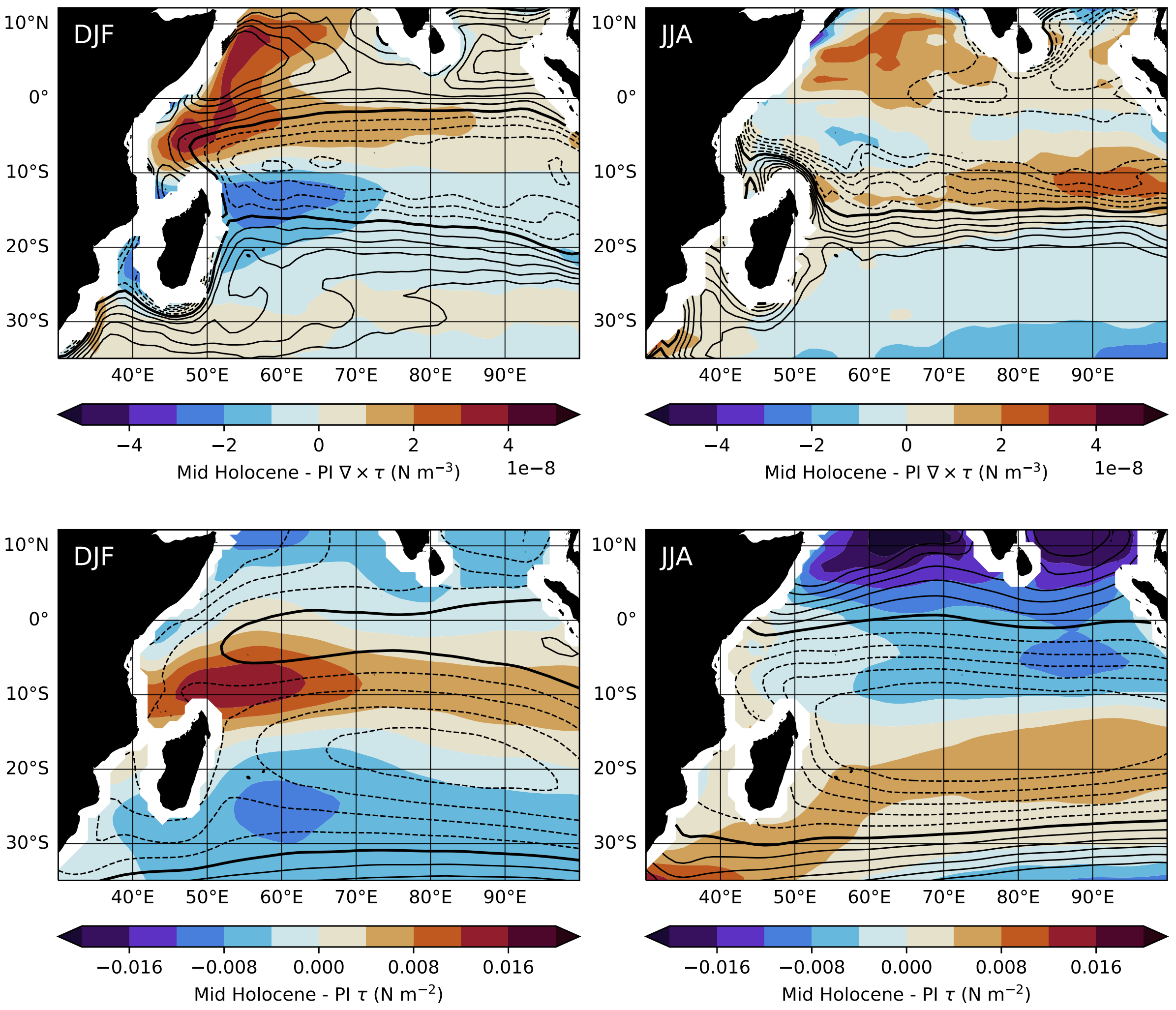


**Supplementary Fig. 15** *Top:* Black contours show the pre-industrial (PI) DJF (left) and JJA (right) wind stress curl, in increments of $2 \times 10^{-8}$ N m$^{-3}$. Dashed lines indicate negative curl. Colours show the mid Holocene − PI wind stress curl (i.e. reds indicate that the wind stress curl was more positive during the mid Holocene). *Bottom:* Black contours show the PI DJF (left) and JJA (right) zonal wind stress, in increments of $2 \times 10^{-2}$ N m$^{-2}$. Dashed lines indicate negative (westward) zonal wind stress. Colours show the mid Holocene − PI zonal wind stress (i.e. reds indicate that the wind stress was more positive, i.e. eastward, during the mid Holocene). These figures are computed from monthly zonal and meridional wind stress at the ocean surface, averaged across contributions from NorESM1-F (Guo et al, 2019), FGOALS-g3 (Zheng et al, 2020), MPI-ESM1-2-LR (Mauritsen et al, 2019), and MIROC-ES2L (Hajima et al, 2020) to PMIP4 (Kageyama et al, 2018).

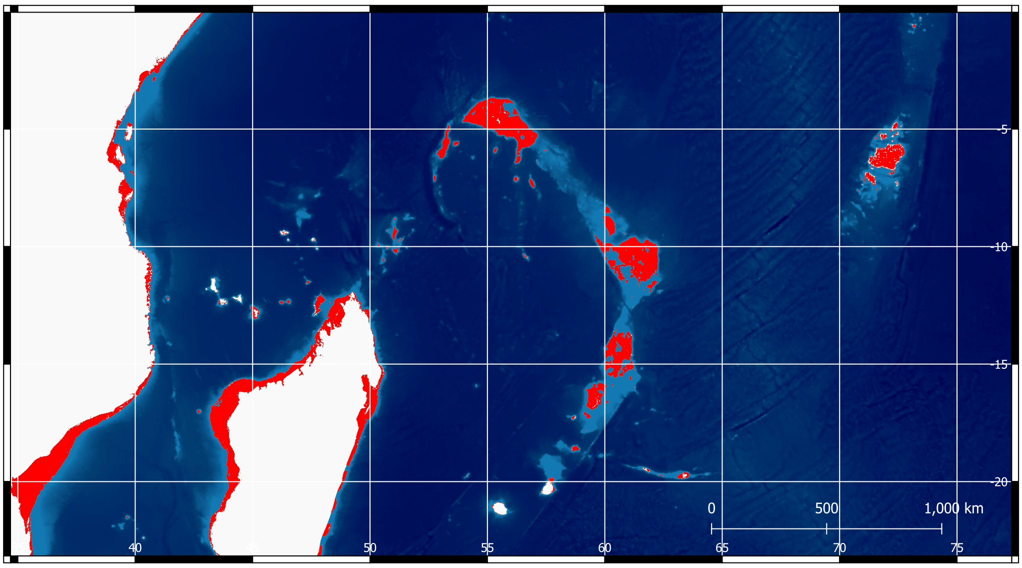


**Supplementary Fig. 16** Exposed land in the present day (white) and under 135 m of eustatic sea-level fall at the last glacial maximum (red), based on bathymetry from GEBCO Compilation Group (2022).

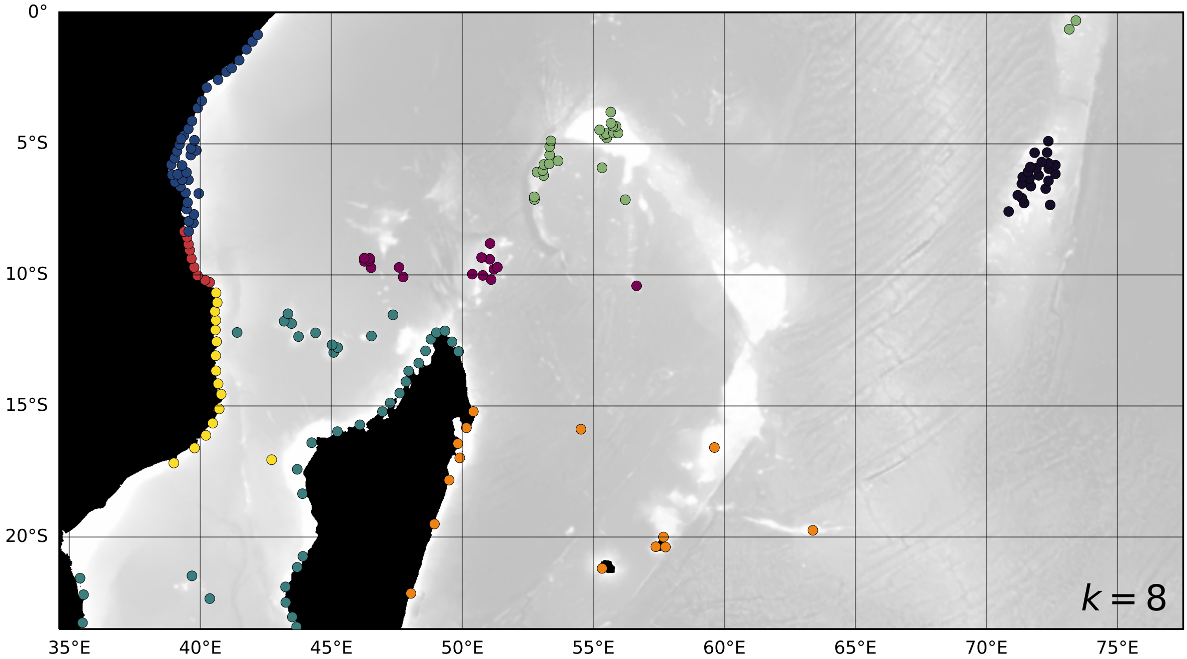


**Supplementary Fig. 17** As in supplementary fig. 8, but with a faster settling rate (6 $d^{-1}$ rather than 1 $d^{-1}$).

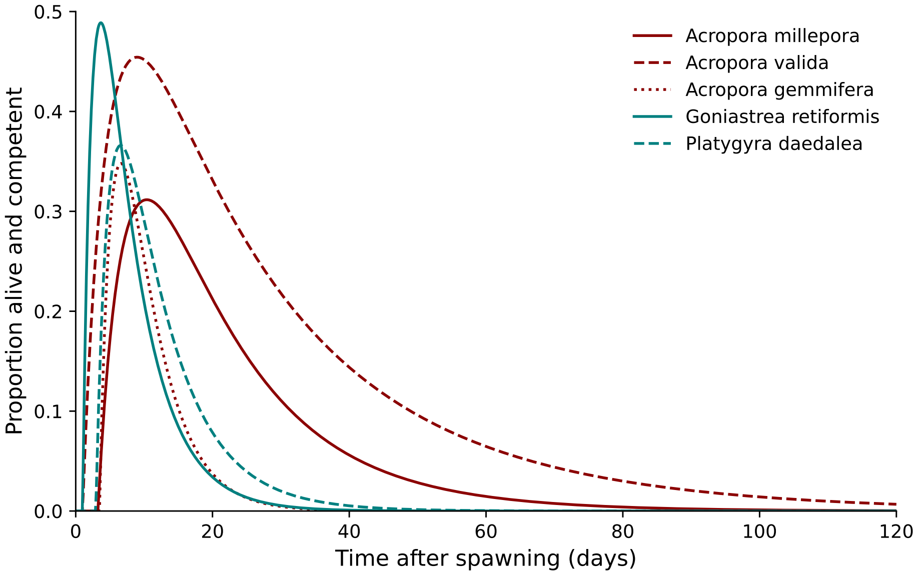


**Supplementary Fig. 18** Proportion of larvae alive and competent as a function of time, for *Platygyra daedalea* and four other coral species, based on observations in Connolly and Baird (2010). As in Vogt-Vincent et al (2023), the minimum allowed competency period is set to 1 day. This diagram is re-used from Vogt-Vincent et al (2023).

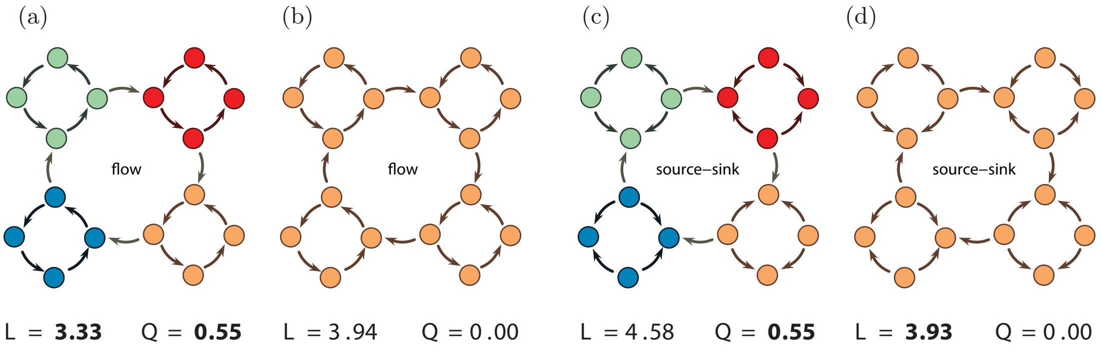


**Supplementary Fig. 19** (a, b) Two possible partitions of network A, and the minimum average code length $L$ (minimised by the map equation) and modularity $Q$ (maximised by modularity). Partition (a) is identified as optimal by both algorithms. (c, d) Two possible partitions of network B, and the minimum average code length $L$ and modularity $Q$. Partition (c) is identified as optimal by modularity, whereas partition (d) is identified as optimal by the map equation. This figure is reproduced with permission from Rosvall et al (2009).
